# Spectral analysis of gene-dependency networks reveals emergent organization across scales in CAR T biology

**DOI:** 10.64898/2026.11.24.768151

**Authors:** Adam Nik, Keenan Hartert

## Abstract

CAR T-cell infusion products are commonly characterized in terms of their cellular populations and transcriptional programs, but these descriptions do not fully capture how a product is organized as a system. We analyzed single-cell RNA sequencing data from the infusion products of 163 patients across eight cohorts to investigate how system-level transcriptional organization relates both to clinical outcome and to patterns of covariation among cellular features across patients. For each patient, we constructed a gene-dependency graph using mutual information between the binary detection states of each gene pair across infusion cells. To characterize each patient’s network architecture, we define spectral organization, *O*_*λ*_, a measure of the relative persistence of collective patterns of gene dependence in the network. Using hierarchical Bayesian survival models, we assessed the associations of both *O*_*λ*_ and various established cellular features with treatment outcome. Higher spectral organization was associated with a higher hazard of treatment failure in both a continuous model (hazard ratio per standard deviation, 1.45, 95% credible interval 1.07–2.19) and a smooth transition model (hazard ratio comparing the high and low regimes, 2.03, 95% credible interval 1.10–3.53). When we investigated how cellular features covary across patients, we found that correlations among both T-cell programs and cell type proportions aligned more coherently with treatment failure risk in the high *O*_*λ*_ regime than in the low regime, revealing a systems-level pattern that helps explain the clinical relevance of spectral organization. Gene set enrichment analysis further showed that more persistent architectures were associated with inflammation and cellular interaction pathways, whereas lower spectral organization was associated with biosynthetic and genome maintenance pathways, consistent with the hypothesis that the organizational state of an infusion product reflects its capacity to reconfigure in response to CAR T treatment. Altogether, our findings identify spectral organization as a property of the infusion product as a whole that connects gene-dependency architecture within individual products to the collective variation of cellular features across patients. This framework therefore motivates studying CAR T biology from a systems perspective alongside established descriptions of cellular state.

**Graphical abstract:** 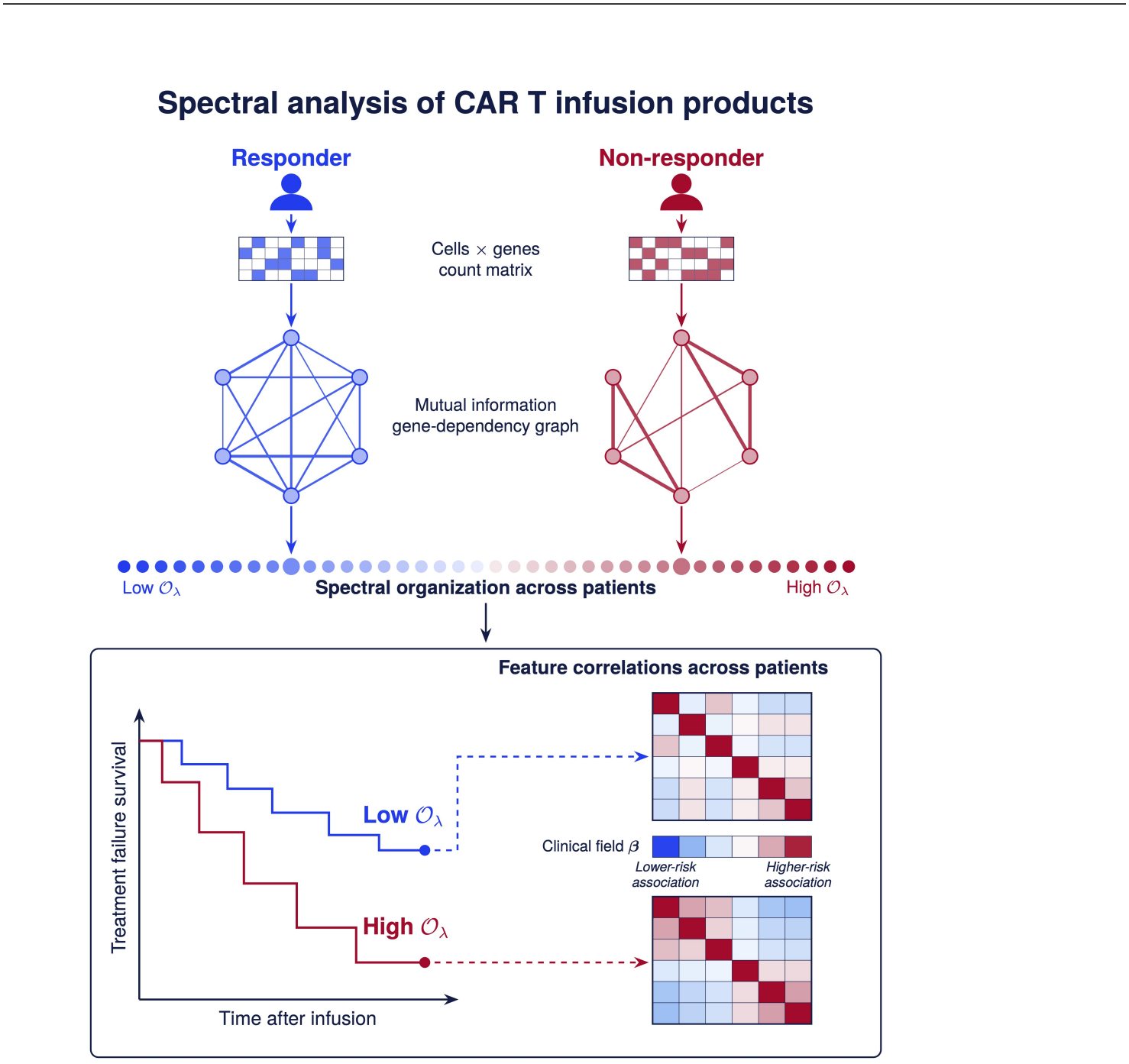

## Introduction

Chimeric antigen receptor (CAR) T-cell therapy is a cancer immunotherapy in which T cells are genetically modified ex vivo to express engineered receptors that better allow them to recognize tumor cells, and then reinfused back into the patient [1, 2]. Despite its clinical success in several hematologic malignancies, many patients still fail to respond or eventually relapse [3, 4]. Treatment outcomes vary across diseases, patient populations, manufactured products, and treatment protocols [5, 6], thus motivating efforts to identify infusion product properties associated with therapeutic response across these clinical contexts [7, 8].

Single-cell RNA sequencing (scRNA-seq) has enabled detailed study of infusion product composition and cellular programs as they relate to patient outcomes [9, 10]. Previous studies have associated naive, stem-like, and memory programs and phenotypes in CAR T-cell infusion products with favorable treatment outcomes [7, 11, 12]. On the other hand, exhaustion-associated programs in infusion products have been linked to poorer responses [7, 11], and regulatory CAR T cells were more abundant in axi-cel infusion products from nonresponders [8]. These prior findings establish cellular state as an important dimension of infusion product characterization to understand determinants of treatment outcome.

However, the strength and consistency of these associations vary across studies. Differences in study design and clinical endpoints further complicate comparisons across cohorts [9, 10]. Furthermore, individual program scores and cell state proportions describe features of an infusion product, but do not directly encode the dependency structure among expressed genes. Products with similar cellular compositions may therefore differ in aspects of organization that these summaries do not capture. This then raises the question of whether infusion product organization is associated with how cellular programs collectively relate to clinical outcome.

Systems and network approaches have provided several ways to study cellular organization across scales as it relates to biological function. Frameworks such as SCENT integrate gene expression with a protein-interaction network to estimate differentiation potency [13], while information-theoretic approaches such as PIDC use dependencies across single cells to infer putative gene-regulatory relationships [14]. In the field of CAR T-cell therapy, Kirouac and colleagues modeled tumor-antigen-regulated transitions among memory, effector, and exhausted states to describe CAR T-cell expansion and tumor response; they identified memory-cell proliferation and effector cytotoxic potency as principal determinants of clinical variation within their model [15]. However, an open question remains as to how system-level properties of an infusion product connect to the collective clinical significance of its cellular components. Here, we ask whether the statistical gene-dependency structure within individual products is associated with how cellular programs collectively relate to clinical risk across patients.

For each patient in this study, we construct a gene mutual information graph using scRNA-seq data from their infusion product. In these graphs, edges represent observed statistical dependencies among genes rather than direct molecular interactions or cell-to-cell communication. We summarize this network using a statistic we coin *spectral organization*, denoted *O*_*λ*_, which we define as a weighted measure of the persistence of nontrivial modes of its normalized adjacency matrix. Persistence here refers to the structural characteristics of the gene statistical network, rather than having a direct biological or physical interpretation.

Evaluating these relationships across studies also requires accounting for differences in clinical populations and follow up schedules. Hierarchical survival models provide an elegant way to pool evidence across studies while allowing associations to vary among them [16]. Furthermore, Bayesian survival models offer approaches for handling partially observed event times and uncertain covariates [17, 18]. In single-cell studies, an additional source of uncertainty arises from the fact that a patient’s cellular composition is estimated from a finite sample of cells [19], and the number of sampled cells can vary substantially between patients. For these reasons, we developed a hierarchical Bayesian survival framework that accounts for finite-cell uncertainty in estimates of spectral organization and cellular features, along with differences in outcome reporting, CAR T product, and disease.

In this study, we analyze infusion product scRNA-seq data across eight CAR T-cell datasets. We find that lower spectral organization was associated with better clinical outcome. Furthermore, while we found that cellular correlation structures were similar across the high and low *O*_*λ*_ regimes, they aligned more closely with a fixed clinical direction in the high *O*_*λ*_ regime. This suggests that favorable and adverse cellular features are best understood in the context of the broader patterns they form, particularly in products with more persistent gene-dependency architectures. Finally, gene set enrichment analysis (GSEA) links higher spectral organization to pathways involved in responding to the cellular environment, such as inflammation and cellular interaction, and lower spectral organization to pathways supporting internal cellular processes, including biosynthesis and genome maintenance. Therefore, our findings connect the transcriptional architecture within a patient to how cellular states collectively relate to clinical risk across patients, supporting the view of CAR T products as organized cellular systems whose properties depend on both their components and the relationships among them.

## Results

### CAR T infusion product data from eight clinical cohorts

We analyzed infusion product single-cell RNA sequencing data from eight publicly available hematologic malignancy CAR T cohorts, spanning both relapsed/refractory B-cell lymphoma and pediatric B-cell acute lymphoblastic leukemia. (Fig. 1; Table 1). The cohorts included axi-cel, tisa-cel, and tandem CD19/CD20 CAR T products. The primary analysis population included 163 patients after restriction to patients with annotated clinical outcomes and at least 1,000 retained cells after preprocessing. Of these patients, there were 105 events and 58 censored. Study-specific outcome definitions and their harmonization are described in the Methods.

**Table 1:** CAR T cohorts included in the primary analysis population. The primary analysis included with patients with both annotated clinical outcomes and at least 1,000 retained infusion product cells after preprocessing. Events and censoring refer to the harmonized treatment failure endpoint.

| Dataset | GEO accession | Disease | CAR T product | Reference | *n* | Events | Censored |
|---|---|---|---|---|---|---|---|
| Bai 2022 | GSE197215 | Pediatric B-ALL | tisa-cel | [12] | 10 | 7 | 3 |
| Bai 2024 | GSE262072 | Pediatric B-ALL | tisa-cel | [20] | 42 | 35 | 7 |
| Cao | GSE207192,<br>GSE283351 | r/r DLBCL | axi-cel | [21] | 9 | 3 | 6 |
| Haradhvala | GSE197268 | r/r B-cell lymphoma,<br>including LBCL | axi-cel,<br>tisa-cel | [8] | 19 | 12 | 7 |
| Li | GSE241783 | r/r LBCL | axi-cel | [22] | 35 | 20 | 15 |
| Maurer | GSE273170 | r/r LBCL | axi-cel | [23] | 4 | 2 | 2 |
| Wang | GSE243325 | r/r DLBCL | Tandem CD19/CD20 | [24] | 6 | 3 | 3 |
| Yu | GSE297676 | r/r LBCL | axi-cel,<br>tisa-cel | [25] | 38 | 23 | 15 |
| **Total** |  |  |  |  | **163** | **105** | **58** |

**Figure 1:**
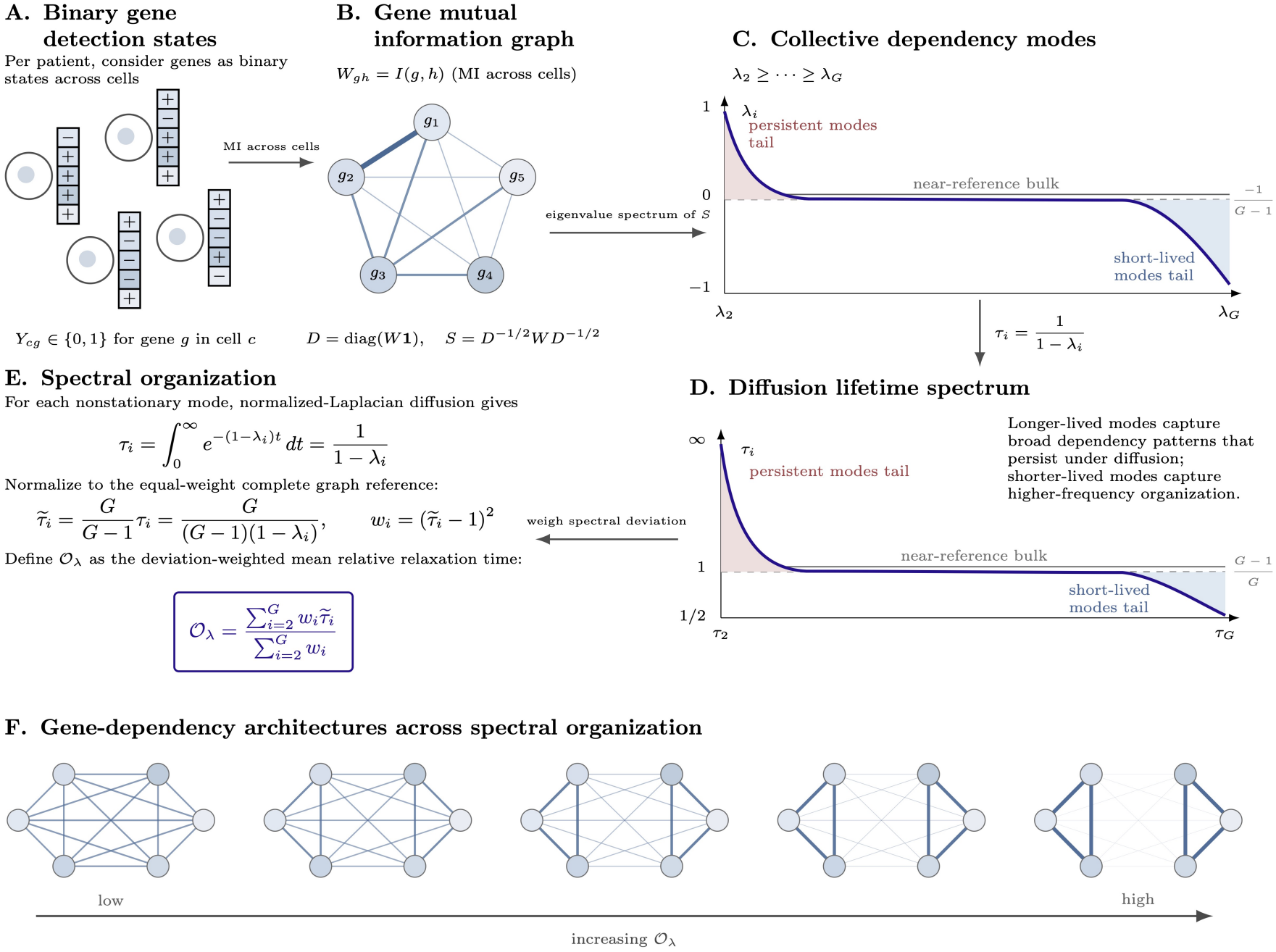
Spectral organization connects single-cell gene detection to patient-level dependency architecture. **A**. For each infusion product, gene detection is recorded as a binary state for each gene in each cell. **B**. For distinct genes *g* and *h*, the edge weight *W*_*gh*_ = *I*(*g, h*) is the mutual information between their detection states across cells. We normalize its adjacency matrix by degree to obtain *S* = *D*^*−*1*/*2^*WD*^*−*1*/*2^. **C**. After excluding the stationary mode, we order the remaining eigenvalues from *λ*_2_ to *λ*_*G*_. **D**. Each eigenvalue defines a normalized-Laplacian diffusion lifetime, *τ*_*i*_ = (1 *− λ*_*i*_)^*−*1^. In panels C and D, red and blue shading mark modes with lifetimes above and below the equal-weight complete-graph reference, respectively. **E**. Relative to this reference, the lifetimes are 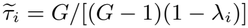. Spectral organization, *O*_*λ*_, averages these relative lifetimes, weighting each mode by its squared departure from the reference, 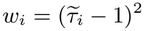. Higher values mean that longer-lived modes account for a greater share of these spectral departures. **F**. Example graphs illustrate increasing *O*_*λ*_ from left to right as edge weights become more concentrated within the two groups of nodes.

### Spectral organization measures the persistence of gene dependencies within an infusion product

We next asked whether the observed statistical dependencies among genes provide a complementary description of infusion product organization. Single-cell expression profiles provide us information to which genes are detected and how their detection states vary together across cells.

We first represented each infusion product as a patient-specific gene-dependency graph (Fig. 1). For a product containing *N* sampled cells and *G* retained genes, let *Y*_*cg*_ ∈{0, 1} indicate whether gene *g* is detected in cell *c*. For genes *g* and *h*, let *I*(*g, h*) denote the mutual information between their binary detection states across cells [26],

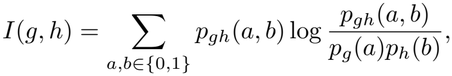

where *p*_*gh*_ denotes the joint detection distribution and *p*_*g*_ and *p*_*h*_ its marginals. We form the symmetric weighted adjacency matrix *W* by setting *W*_*gh*_ = *I*(*g, h*) for *g* ≠*h* and *W*_*gg*_ = 0. Hence, each node represents a gene, and each edge represents the statistical dependency of those genes across sampled cells (these edges do not identify direct molecular interactions nor cell-to-cell communication). Furthermore, note that the construction of *W* requires no predefined cell labels or transcriptional programs. Additional details of graph construction and spectral calculations are provided in the Methods.

To characterize dependency architecture, define the symmetric normalized adjacency matrix as

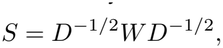

where *D* = diag(*W* **1**) contains the weighted node degrees. For a connected graph with *G*≥ 2, the eigenvalues of *S* satisfy [27]

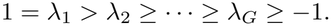

Here, graph diffusion describes how patterns of variation relax across the gene-dependency network. The normalized Laplacian *L*_sym_ = *I* − *S* governs this, with each eigenvalue of *S* specifying the decay rate of a corresponding collective mode [27]. The mode corresponding to *λ*_1_ = 1 is stationary and is excluded from the summary. Each remaining mode decays exponentially at rate *ν*_*i*_ = 1 − *λ*_*i*_ *>* 0, giving the diffusion lifetime

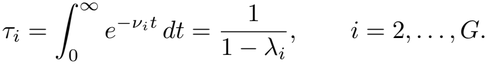

Thus, smaller nonzero Laplacian eigenvalues correspond to longer diffusion lifetimes, or lower graph frequencies [28].

Because *W* has zero diagonal, tr(*S*) = 0. Together with *λ*_1_ = 1, this gives us the exact mean of the nonstationary eigenvalues,

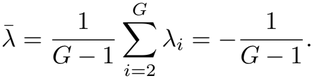

Note that this is also the eigenvalue shared by all nonstationary modes of an equal-weight complete graph. We therefore use that graph as a uniform-connectivity reference, with lifetime

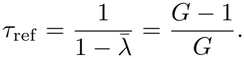

Normalizing each lifetime by this reference gives us

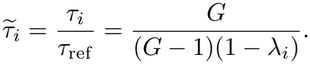

Hence, 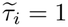 denotes the complete-graph reference for any retained gene count.

We define the deviation-weighted mean relative relaxation time as

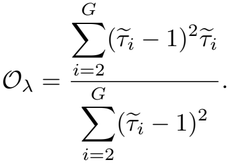

We refer to this quantity as *spectral organization* (Fig. 1). This statistic averages relative diffusion lifetimes, weighting each nonstationary mode by its squared deviation from the uniform, complete reference. Thus, higher *O*_*λ*_ values describe patient dependency architectures of greater spectral persistence, dominated by lower frequency modes. Note that for a uniform, complete graph, the ratio is undefined; thus, we assign *O*_*λ*_ = 1 by continuous extension (Supplement).

This weighting distinguishes spectral organization from the spectral gap and the normalized Kemeny’s constant. The spectral gap, 1 − *λ*_2_, is the decay rate of the slowest nonstationary mode [27]. Kemeny’s constant sums the diffusion lifetimes of all nonstationary modes [29]. Dividing by its value (*G* − 1)^2^*/G* for the equal-weight complete graph gives

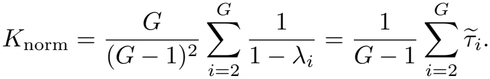

Thus, *K*_norm_ gives equal weight to all relative diffusion lifetimes, while *O*_*λ*_ gives more weight to modes whose lifetimes depart further from the reference. We compared all three summaries under the same patient-bootstrap survival analysis. The continuous association with treatment failure had a 95% credible interval excluding one for *O*_*λ*_ alone (Supplementary Fig. S2; Supplementary analysis).

### Estimating spectral organization from finite cell samples

Any finite sample estimation of mutual information is biased; in other words, the expected value of its estimation is dependent on the number of samples used during estimation [30]. Because *O*_*λ*_ is a nonlinear function of the mutual information graph *W*, its estimation can also be biased and vary with sampling depth. To accommodate this, for each patient, we computed *O*_*λ*_ across cell subsamples of various cell counts, spanning 50 cells to all cells available, where we reconstructed the mutual information graph and subsequently assessed *O*_*λ*_ at each depth. This produced a patient-specific trajectory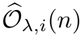 that describes how the estimated dependency architecture changes with sampling depth. Thus, our target estimand for each patient is the limiting value of their trajectory as sampling depth increases, denoted 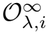 for patient *i*.

To estimate 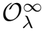 as the asymptote of this trajectory, we used a hierarchical empirical-Bayes model with patient-specific asymptote, slope, curvature, and convergence timescale. The model was fitted on the log-excess scale,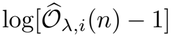.^1^ The uncertainty around the central estimate 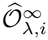 reflects the information available in patient *i*’s observed trajectory. Validation using truncated trajectories is presented in Supplementary Fig. S1. We refer to 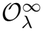 as simply *O*_*λ*_ for the remainder of the paper unless otherwise specified.

### Higher spectral organization is associated with higher hazard of treatment failure

To evaluate clinical associations across heterogeneous cohorts, we used a hierarchical Bayesian survival model that incorporated exactly observed, interval-censored, and right-censored outcomes (Methods). Each cohort contributed the event timing and follow up information available from its clinical annotations. For a given feature, the model estimated a population-level association with dataset-specific deviations, where we borrowed information across cohorts through partial pooling. Here, dataset-specific baseline hazards allowed underlying event rates to differ, while CAR T product and disease type were included as clinical covariates.

We evaluated each biological feature in a separate model and reported hazard ratios per global standard deviation increase across the complete analysis population. Patient-specific uncertainty was propagated through the survival likelihood by integrating over discrete representations of each feature’s estimated distribution. We assessed both the magnitude of the population-level association and its directional consistency across cohorts.

Let *P* (*β >* 0) denote the posterior probability of a positive population-level coefficient. Higher spectral organization was associated with a higher hazard of treatment failure (HR 1.45 per global standard deviation increase, 95% CrI 1.07–2.19, *P* (*β >* 0) = 0.989). The posterior probability that all eight cohort effects shared a direction was 0.800 within the partially pooled model. Hence, the results support a shared adverse association across the cohorts.

### Higher spectral organization marks a clinically adverse regime

The observation of more adverse outcomes at higher values of *O*_*λ*_ motivated a complementary analysis of low vs. high spectral organization regimes. We wanted a binary analogue of the continuous survival model that would assess differences in treatment failure outcome between the low and high spectral organization regimes, while allowing group membership to be learned probabilistically rather than fixed by any predefined threshold. As a result, we extended the hierarchical Bayesian survival framework to a smooth transition model whose location was inferred from clinical outcomes (Methods). The model estimated the high-versus-low hazard contrast with the baseline hazard defined at the transition midpoint. The model retained the same dataset-specific baseline hazards, partially pooled dataset effects, adjustment for CAR T product and disease, and integration over patient-specific uncertainty in *O*_*λ*_ as in the continuous survival model. Similarly, we estimated the crossover on the common absolute spectral organization scale, restricting its location to the 10^th^–90^th^ percentile range of the analysis population. Thus, each patient’s inferred degree of membership in the respective regimes reflected uncertainty from both their *O*_*λ*_ value and the estimated crossover location.

The resulting population-level hazard ratio between the high and low *O*_*λ*_ regimes was 2.03 (95% CrI 1.10– 3.53, *P* (*β >* 0) = 0.986). The posterior probability that all eight dataset-specific regime effects were adverse was 0.800. As in the continuous analysis, this probability summarizes directional agreement within the partially pooled model. The inferred crossover point was centered at *O*_*λ*_ = 1.206 (95% CrI 1.112–1.457), which marks the value of *O*_*λ*_ at which the fitted log hazard changed most rapidly.

Therefore, higher spectral organization was associated with greater treatment failure hazard under both the continuous and smooth transition models. For visualization purposes, we grouped patients into low vs. high spectral organization regime groups according to whether their *O*_*λ*_ estimate fell below or above the posterior-median transition point, and then estimated pooled survival curves using their interval-censored event times (Fig. 2).

**Figure 2:**
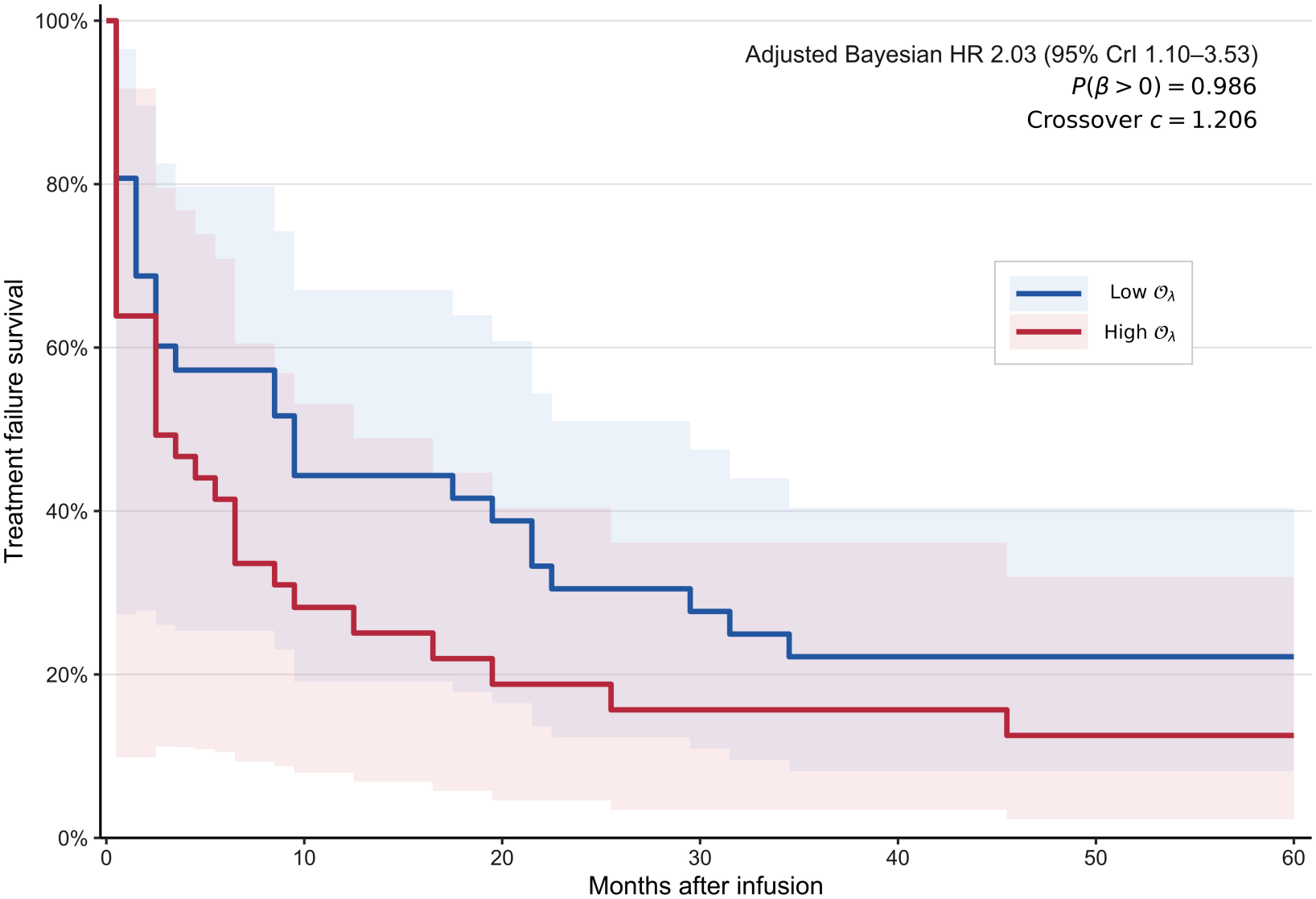
Treatment failure survival across spectral organization regimes. Patients with central *O*_*λ*_ estimates below the posterior-median crossover (*c* = 1.206) were grouped as low (blue). Those at or above it were grouped as high (red). Curves are pooled, unadjusted Turnbull estimates of interval-censored survival, retaining reported event intervals and right-censoring patients without an observed event at their last follow up. Shading shows 95% confidence intervals calculated on the log–log scale for the displayed groups; these intervals exclude uncertainty in spectral estimates and the crossover. The annotated HR of 2.03 (95% CrI 1.10–3.53, *P* (*β >* 0) = 0.986) compares the limiting high and low hazards in the midpoint-centered Bayesian model. That model incorporates predictor uncertainty, dataset-specific baseline hazards and effects, and treatment and disease adjustment. The curves visualize the association in sample, because the crossover was learned from the same clinical outcomes.

### The association with treatment outcome extends to held-out cohorts

To assess whether the association depended on any single cohort, we refitted both survival models after omitting each dataset in turn. Posterior-median hazard ratios remained above one in every seven-cohort refit, and the 95% credible intervals excluded one in six of eight continuous and five of eight smooth transition fits (Supplementary Table S2). Thus, the adverse direction of *O*_*λ*_ persisted under every cohort omission.

**Supplementary Fig. S1:**
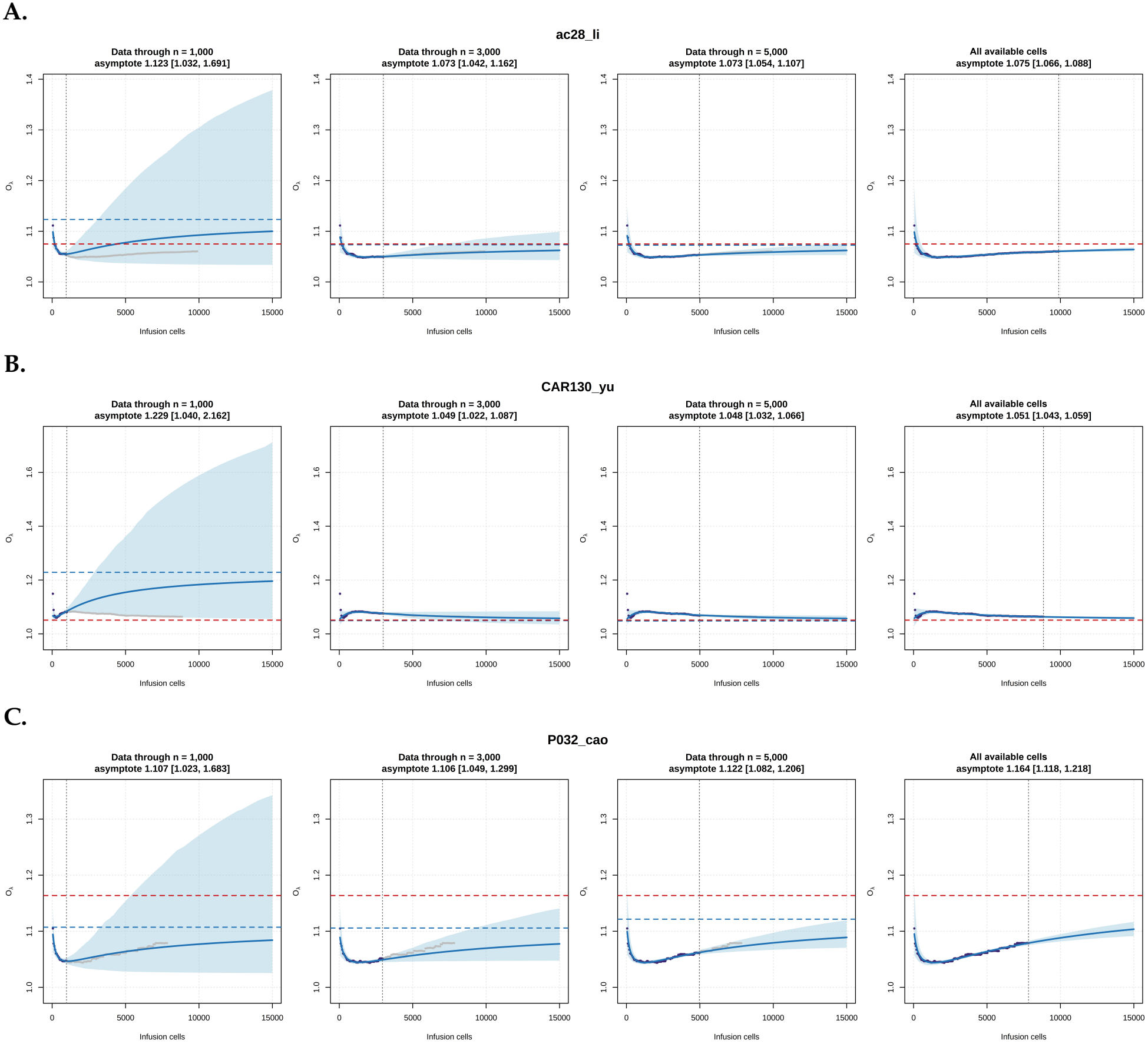
Representative finite-depth validation trajectories. Finite-depth inference in three representative held-out infusion products: (**A**) ac28_li, (**B**) CAR130_yu, and (**C**) P032_cao. Columns use observations through 1,000, 3,000, 5,000, or all available cells. Panels share a *y*-axis within rows and extend to 15,000 cells. Purple and gray points show fitted and later withheld observations, respectively. Blue curves and light-blue bands show inferred latent trajectories and pointwise 95% credible intervals; the bands do not represent prediction intervals for new observations. Dashed lines show the central asymptote estimate 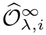 fitted from each prefix (blue) and the complete trajectory (red). Vertical dotted lines mark the maximum fitted depth. Only the red asymptote reference appears in complete-trajectory panels. Examples show stabilization at moderate depth (**A**), narrowing of early uncertainty (**B**), and refinement of the extrapolated asymptote along a gradually rising trajectory (**C**). Curves beyond observed depths are extrapolated, and the complete-trajectory reference is itself inferred.

**Supplementary Table S1:** Finite-depth validation of inferred asymptotic spectral organization. Leave-one-patient-out validation included 16 patients with at least 7,500 infusion product cells. Asymptote MAE, RMSE, and Spearman correlation compare prefix and complete-trajectory estimates fitted under the same held-out hierarchy. Reference containment reports the number of finite-depth 95% intervals containing the corresponding central complete-trajectory estimate, out of 16. Withheld-trajectory MAE is the mean of patient-level MAEs. Errors and interval widths use the *O*_*λ*_ scale.

| Maximum<br>depth | Asymptote<br>MAE | RMSE | Spearman<br>*ρ* | Reference<br>containment | Median 95%<br>interval width | Withheld<br>MAE |
|---|---|---|---|---|---|---|
| 1,000 | 0.1245 | 0.1749 | 0.31 | 16/16 | 0.793 | 0.0292 |
| 3,000 | 0.1019 | 0.2105 | 0.80 | 13/16 | 0.126 | 0.0169 |
| 5,000 | 0.0620 | 0.1282 | 0.94 | 12/16 | 0.061 | 0.0101 |

Next, we evaluated whether associations learned from the remaining cohorts extended to patients in an omitted dataset. For each patient, we constructed cross-fitted risk scores using models trained without that patient’s dataset, with scaling and transition parameters determined from the training data (Supplement). In the pooled evaluation of held-out patients, a one standard deviation increase in the cross-fitted score was associated with higher treatment failure hazard for both the continuous score (HR 1.50, 95% CrI 1.04–2.42) and the transition score (HR 1.84, 1.17–2.93) (Supplementary Table S3). As so, the association with treatment failure persisted across all cohort omissions, and spectral organization remained prognostic in patients from unseen cohorts.

### Cell resampling preserves the higher hazard of treatment failure

Finally, we tested whether the survival association depended the on asymptotic trajectory inference method. For each patient, we generated 25 cell resamples using Bayesian-bootstrap weights, and then reconstructed the gene-dependency graph and computed *O*_*λ*_ for each resample. We tested the resulting *O*_*λ*_ estimates through the same adjusted survival models. The continuous association remained under direct resampling (HR 1.51 per standard deviation, 95% CrI 1.12–2.08). Moreover, despite the narrower spread of finite-cell bootstrap estimates across patients (interquartile range 0.049, compared with 0.186 for the asymptotic estimates), the transition model still yielded a high-versus-low HR of 1.80 (95% CrI 0.97–3.19, *P* (*β >* 0) = 0.969) (Supplementary Table S6; Supplementary analysis). Thus, direct graph reconstruction retained the adverse association, with greater uncertainty about the magnitude of the regime contrast. We retained 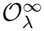 as the primary estimand because it targets a common infinite-depth limit, whereas bootstrapping to the same cell count still depends on how many cells were originally observed for each patient.

### Transcriptional programs and cellular phenotypes across infusion products

Once we established the clinical importance of spectral organization, we then moved onto studying its relationship to established T-cell biology. To do this, we characterized the infusion products using two complementary representations; one at the level of transcriptional programs, quantified by T-CellAnnoTator (TCAT) [31], and the other at the level of cellular phenotypes, labeled using the open Functional CAR-T Atlas, [32].

TCAT represents each cell’s gene expression profile as a weighted combination of shared gene expression programs (GEPs) derived by nonnegative matrix factorization (NMF). Additionally, TCAT computes activation and proliferation scores, which each summarize a prespecified combination of these programs (Methods).

While TCAT decomposes a cell’s gene count vector into continuous scores, we also sought to label each cell into known T-cell types. In order to do so, we mapped infusion cells to the atlas’s 11 annotated phenotypes using an atlas-derived scANVI model, which was trained on a gene panel shared across our datasets. These annotations included CD4 subtypes such as central memory, effector memory, and cytotoxic states; CD8 subtypes such as memory, effector memory, and cytotoxic states; and other labels including regulatory, proliferative, apoptotic, ribosomal-enriched, and monocyte-like T-cell phenotypes. Each analyzed cell was assigned its most likely atlas label from the scANVI model (Methods).

For each infusion product, we calculated mean GEP usages and atlas phenotype proportions. Since mean GEP usages and phenotype proportions each sum to one within a patient, we treated these as separate compositions and applied a centered log-ratio (CLR) transformation, which expresses each feature relative to the geometric mean of its corresponding composition. Antigen-specific activation (ASA) and proliferation scores were computed from their corresponding CLR-transformed GEPs. Finally, to represent uncertainty from sampling a finite number of cells, we repeatedly assigned random Bayesian-bootstrap weights to cells and recalculated the patient-level features. Each resulting particle contained a jointly computed set of features, preserving their dependence and compositional constraints (Methods).

### Survival analysis of cellular programs and phenotypes recovers established treatment outcome associations

We evaluated the clinical associations of the 52 TCAT gene expression programs, TCAT ASA and proliferation composite scores, and the 11 atlas-defined phenotype proportions across all 163 patients. Each feature was evaluated separately using the hierarchical survival framework, adjusting for CAR T product and disease, allowing associations to vary by dataset, and accounting for finite-cell uncertainty. In the continuous models, population-level 95% credible intervals (CrIs) excluded a hazard ratio of 1 in eight GEPs, one score, and four atlas phenotypes (Fig. 3A,B,D). Again, HRs describe a global standard deviation increase in each transformed feature.

**Figure 3:**
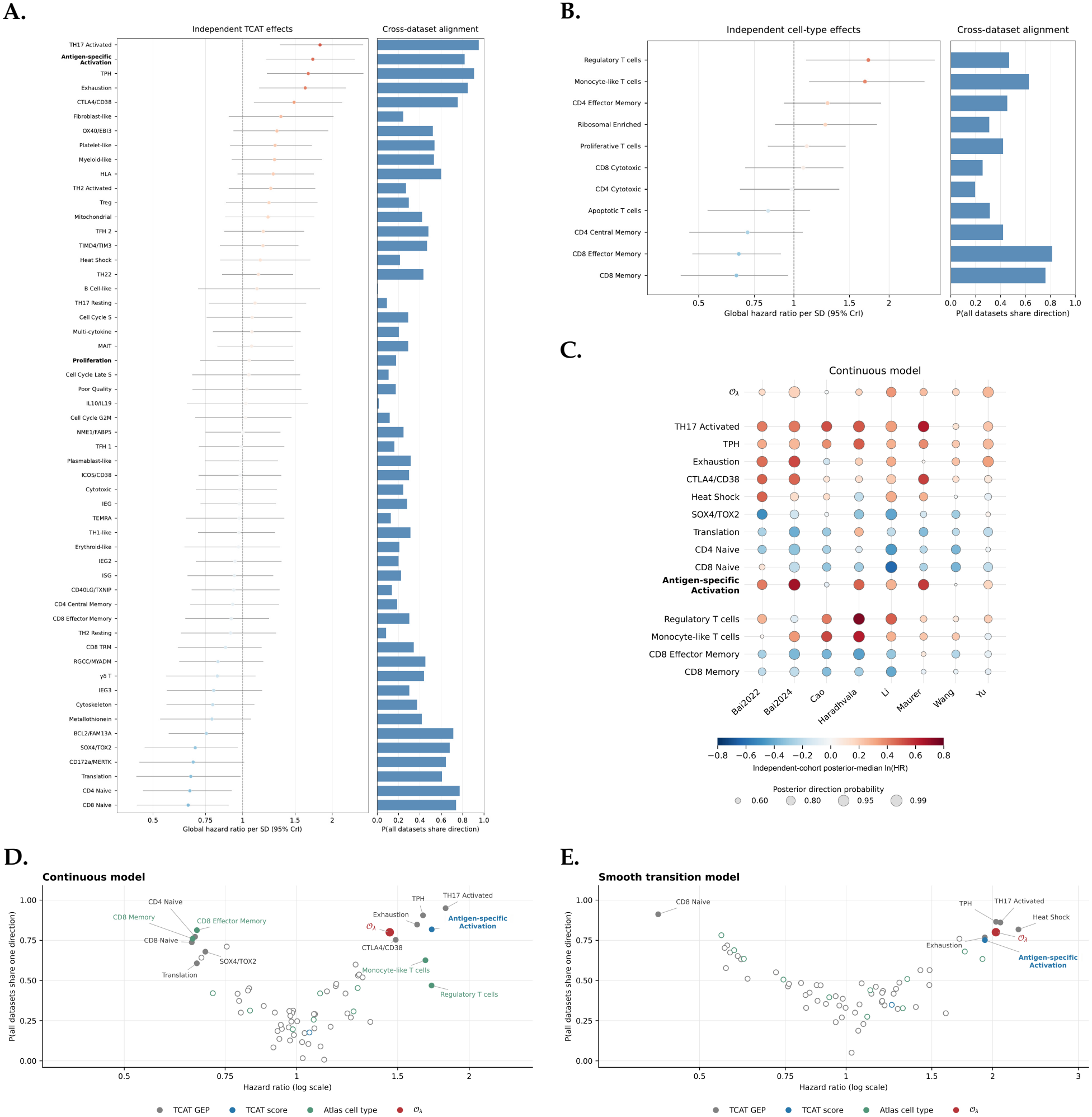
Outcome associations of infusion product programs, phenotypes, and spectral organization across eight cohorts. **A, B**. Continuous-model associations for 52 TCAT GEPs and two composite scores (**A**) and 11 atlas phenotype proportions (**B**), fitted individually. Points show population-level posterior median HRs per global standard deviation, with 95% CrIs. Bars show the posterior probability that all cohort effects share a direction. **C**. Independent continuous-model fits in each cohort for predictors whose population-level interval excluded one in either formulation. Color shows posterior-median ln(HR) (red, higher risk; blue, lower risk); point size shows posterior support for the estimated direction. **D, E**. Population-level HRs and posterior probabilities of a shared direction for continuous (**D**) and smooth transition (**E**) models. Each point represents a predictor, with filled points denoting 95% CrIs excluding one. HRs are plotted on logarithmic axes. Smooth transition HRs compare the limiting high and low hazards in the midpoint-centered model. Full cohort models include treatment and disease adjustment and account for dataset heterogeneity.

The survival results captured many of the established findings of the field. CD8 naive and CD4 naive programs were associated with lower event risk (HR 0.66, 95% CrI 0.44–0.90, and 0.66, 0.46–0.92, respectively). Dysfunction-related programs also showed adverse associations, including exhaustion (1.62, 1.14–2.22) and CTLA4/CD38 (1.49, 1.09–2.16). Additionally, several activation and related programs showed adverse associations, including activated Th17 (1.82, 1.33–2.54), the antigen-specific activation composite score (1.72, 1.20–2.38), and T peripheral helper (Tph; 1.66, 1.21–2.54). The proliferation composite score (1.05, 0.72–1.49) and cytotoxic GEP (0.98, 0.70–1.30) had no clearly resolved individual association.

The atlas cell types showed a similar pattern. Higher relative abundance of CD8 memory and CD8 effector memory phenotypes was associated with lower event risk (HR 0.66, 0.44–0.96, and 0.67, 0.48–0.91, respectively). On the other hand, regulatory T-cell and monocyte-like phenotypes were associated with higher risk (1.72, 1.09–2.79, and 1.68, 1.12–2.59, respectively). That being said, phenotype abundance and program usage still captured slightly different aspects of the product. For example, the favorable association of the CD8 effector memory phenotype was less apparent in the similarly named GEP (0.92, 0.64–1.23).

In the smooth transition models, the high regime was associated with higher hazard for activated Th17 (HR 2.07, 95% CrI 1.21–3.63), Tph (2.03, 1.17–3.49), exhaustion (1.93, 1.003–3.57), heat shock (2.26, 1.02–4.25), and ASA (1.93, 1.06–3.39) (Fig. 3E). The CD8 naive GEP showed the clearest association with lower hazard (HR 0.41, 0.24–0.76). The estimate for the CD4 naive GEP was also below one, though its interval included one (0.58, 0.34–1.01). Among the atlas phenotypes, median hazard ratios remained below one for CD8 memory and CD8 effector memory and above one for regulatory and monocyte-like T cells. The credible intervals for all 11 atlas contrasts included one.

Some features showed strong associations to treatment outcome through their hazard ratios, yet have wavering levels of dataset agreement. For example, the TCAT ASA score and the atlas regulatory T-cell proportion both had continuous model HRs that rounded to 1.72; however, the posterior probability that all eight dataset effects had the same sign was 0.818 for ASA and 0.469 for the regulatory T-cell proportion (Fig. 3D). Several features also did show strong cross-cohort directional support. This probability was 0.949 for the activated Th17 GEP and 0.905 for Tph GEP, and 0.813 for the atlas CD8 effector-memory phenotype. For comparison, the corresponding probability for spectral organization was roughly 0.800 for both the continuous model and transition models.

To further assess the heterogeneity in treatment failure associations between cohorts, we fit separate continuous models for each cohort for the 15 predictors whose population-level 95% CrI excluded one in either the continuous or smooth transition model (Fig. 3C; Methods). The heat shock program showed the most variation in direction, with median HRs above one in five cohorts and below one in three. The magnitude of the estimated associations also varied across cohorts; for example, the median HR for antigen-specific activation was 0.93 in the Cao cohort and 1.95 in Bai 2024 cohort. Similarly, the median HR for regulatory T-cell abundance ranged from 0.91 in Bai 2024 to 2.11 in Haradhvala. Several predictors did show a consistent adverse direction, as median HRs exceeded one in all eight cohorts for activated Th17, Tph, and CTLA4/CD38. For *O*_*λ*_, median HRs exceeded one in seven cohorts, with the Cao estimate close to one (HR 0.98, 95% CrI 0.42–2.30).

Finally, we evaluated spectral organization in two joint models, one including all TCAT GEPs and the other including all atlas cell type proportions, to examine its conditional association with outcome (Methods). With all 52 GEPs included, the continuous HR for *O*_*λ*_ was 1.17 (0.85–1.74, *P* (*β >* 0) = 0.831). With all 11 atlas proportions, it was 1.19 (0.96–1.60, *P* (*β >* 0) = 0.941). For the joint transition models, we held each crossover fixed at its individual-model posterior median. The high-versus-low *O*_*λ*_ regime HR was 1.65 (0.90–3.34, *P* (*β >* 0) = 0.946) alongside the GEPs and 1.59 (0.97–2.94, *P* (*β >* 0) = 0.962) alongside atlas composition. The reduction in posterior support extended to cellular features. Among TCAT GEPs, the number of continuous associations whose 95% CrIs excluded one fell from eight in the individual feature models to just one in the joint model. Among atlas phenotypes, it fell from four to zero. The corresponding count for GEP transition contrasts fell from five to two, with CD8 naive and RGCC/MYADM retaining intervals that excluded one. Similarly, all atlas transition intervals included one in both the individual and joint models. This is not surprising as joint models estimate conditional associations among correlated features under regularization, making individual contributions more difficult to resolve.

Our survival analysis recovered several established patterns in CAR T infusion products. Higher usages of naive T-cell programs were associated with lower treatment failure hazard, while exhaustion and several activation-associated programs were associated with worse outcomes. Spectral organization showed a broadly consistent adverse direction across cohorts and joint models. We next asked how cellular programs and phenotypes covary across patients and whether their relationships align differently with clinical risk across spectral regimes.

### TCAT programs and atlas cell types covary but only partly capture spectral organization

We next moved to studying how the *O*_*λ*_, TCAT GEPs/scores, and atlas phenotypes all covaried with one another. Using the full patient population, we estimated dataset-adjusted Bayesian-bootstrap correlations among the 52 TCAT gene expression programs (GEPs), 11 atlas-defined T-cell phenotypes, and between each GEP and each phenotype. We also related each feature to continuous *O*_*λ*_. Each draw incorporated uncertainty in the patient-level estimates and Bayesian-bootstrap weights across patients. Features were centered within each dataset and draw before correlations were calculated (Methods).

The GEP correlation matrix shows groups of programs with related functions (Fig. 4A). The three cell cycle programs covaried strongly (posterior-median *r* = 0.79–0.92), as did the CD4 naive and CD8 naive programs (*r* = 0.87). Furthermore, CD8 effector memory usage was linked to cytotoxicity (*r* = 0.86) and to the *γδ* T-cell program (*r* = 0.78). Along the same lines, the atlas recovered recognizable patterns in the cell composition space (Fig. 4B). For example, products relatively enriched for CD8 effector memory cells also tended to be enriched for CD8 memory cells (*r* = 0.60) and depleted of proliferative T cells (*r* = −0.64).

**Figure 4:**
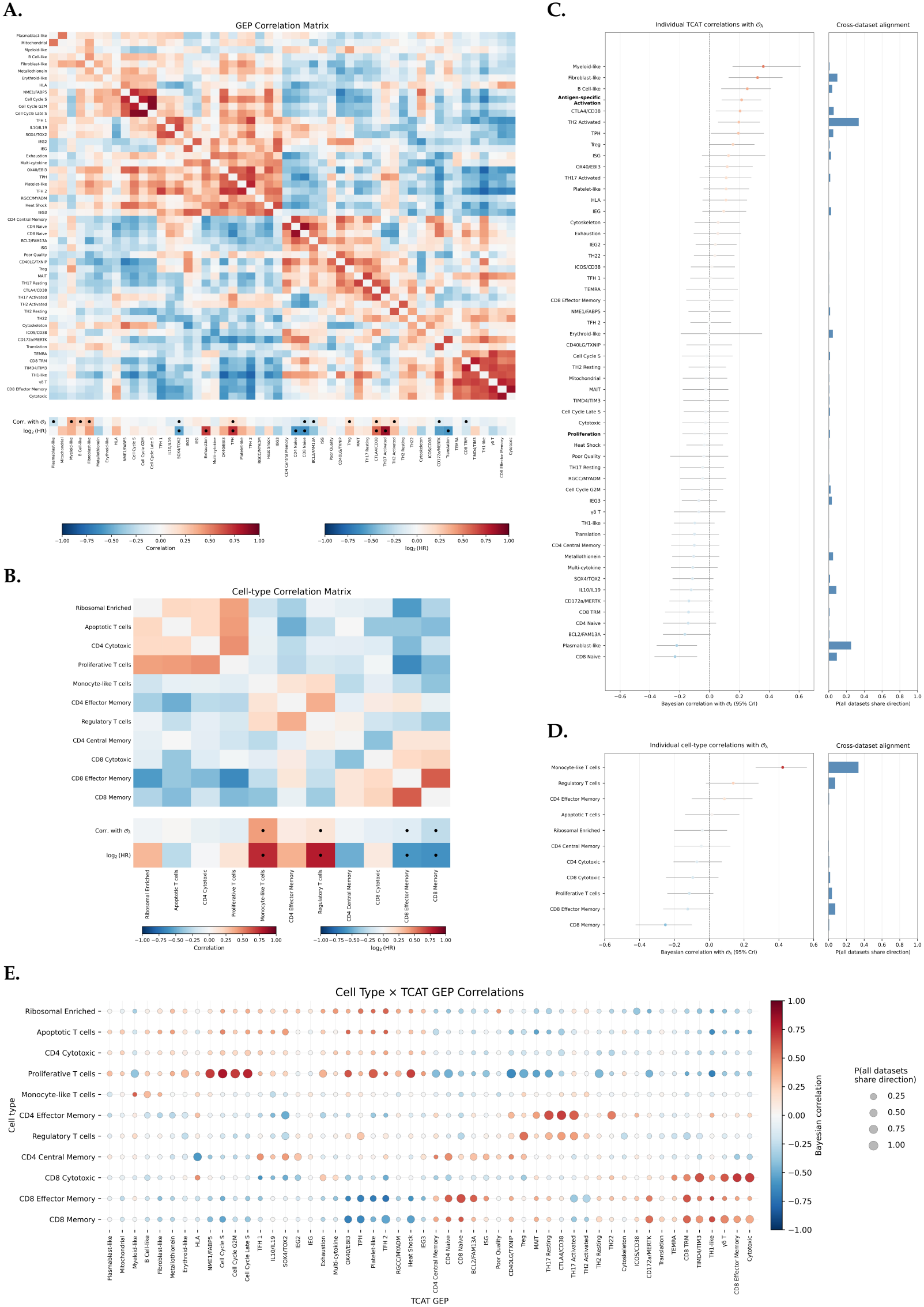
Correlation structure of CAR T infusion products across cellular representations. **A, B**. Dataset-adjusted correlations among TCAT gene expression programs (GEPs) and among atlas cell type proportions. Aligned rows show each feature’s correlation with asymptotic spectral organization and its continuous-model log_2_(HR). Black markers indicate posterior support of at least 0.95 for the correlation direction or a 95% HR credible interval excluding one in the respective rows. **C, D**. Correlations between *O*_*λ*_ and TCAT GEPs and composite scores (**C**) or atlas cell type proportions (**D**), with 95% credible intervals and posterior support for a shared direction across datasets. **E**. Correlations between atlas cell type proportions and TCAT GEP usages, colored by posterior median and sized by the posterior probability of a shared direction across datasets.

Direct comparison of GEP usages with atlas cell type proportions showed where the two representations converged (Fig. 4E). The myeloid-like GEP and monocyte-like T-cell proportion correlated with each other (*r* = 0.60, 95% CrI 0.49–0.69). Correspondence also extended to established T-cell functions; the atlas CD8 cytotoxic proportion correlated with the TCAT cytotoxic and CD8 effector memory programs (*r* = 0.69 and 0.71, respectively). The clearest example actually came from proliferation; the atlas proliferative T-cell proportion correlated with the S-phase (*r* = 0.81), late S-phase (*r* = 0.76), and G2M programs (*r* = 0.70). For each of these three cell-cycle pairs, the posterior probability that all eight datasets shared a correlation direction exceeded 0.96.

Compared to the relatively strong feature-to-feature correlations, individual feature associations with *O*_*λ*_ were more modest (Fig. 4C,D). Of the 52 GEPs, 46 had an absolute posterior-median correlation below 0.20. The strongest positive correlates between the two cellular feature representations were the myeloid-like mixed-lineage GEP (*r* = 0.36, 95% CrI 0.15–0.61) and the atlas monocyte-like T-cell proportion (*r* = 0.42, 95% CrI 0.27–0.56). However, other mixed-lineage-associated signals varied in both directions with spectral organization; fibroblast-like and B-cell-like program signals increased with *O*_*λ*_ (*r* = 0.32 and 0.25, respectively) whereas the plasmablast-like signal decreased (*r* = −0.22). Higher *O*_*λ*_ also accompanied lower CD8 naive program usage (*r* = 0.23, 95% CrI −0.37 to −0.08) and lower relative abundance of atlas CD8 memory cells (*r* = −0.25, 95% CrI −0.42 to −0.10). Among the composite scores, antigen-specific activation was modestly associated with spectral organization (*r* = 0.21, 95% CrI 0.08–0.35), whereas proliferation showed little association (*r* = −0.03, 95% CrI −0.18–0.12).

These correlation analysis findings show that spectral organization is related to, but not recovered by, any individual transcriptional program or cellular phenotype. Even though mixed-lineage-associated features showed the strongest correlations, no single feature closely tracked *O*_*λ*_, and most associations were modest. Finally, we asked whether combinations of cellular features could reconstruct spectral organization. We evaluated four regression approaches for reconstructing asymptotic *O*_*λ*_ in held-out patients from datasets represented in training. The best reconstruction achieved *R*^2^ = 0.313 with TCAT GEPs and *R*^2^ = 0.216 with atlas cell type proportions (Supplementary Table S7). We also constructed a synthetic product for each patient by sampling cells from other patients in the same dataset to match the target patient’s atlas cell type composition. The ranking by asymptotic *O*_*λ*_ agreed only partly between synthetic and observed products (Spearman *ρ* = 0.53, Supplementary analysis). Ultimately, these results suggest that cellular programs and cell type composition capture only an incomplete picture of the meaningful variation of *O*_*λ*_ across patients.

### Spectral organization distinguishes how programs and cell types align with clinical risk

The whole-network construction of *O*_*λ*_ led us to ask whether spectral organization distinguishes how cellular features vary together relative to their clinical associations (Fig. 5A). For each of the two feature representations, TCAT gene expression programs and atlas cell type proportions, let the *clinical-association field* be defined as the vector ***β*** whose entry *β*_*j*_ is the logarithm of the population level posterior median hazard ratio for feature *j* from its individual continuous survival model. Using this field as a common clinical reference, we compared feature correlations between the low and high spectral regimes, with the purpose of studying how relationships among features align with the clinical field in the two regimes.

**Figure 5:**
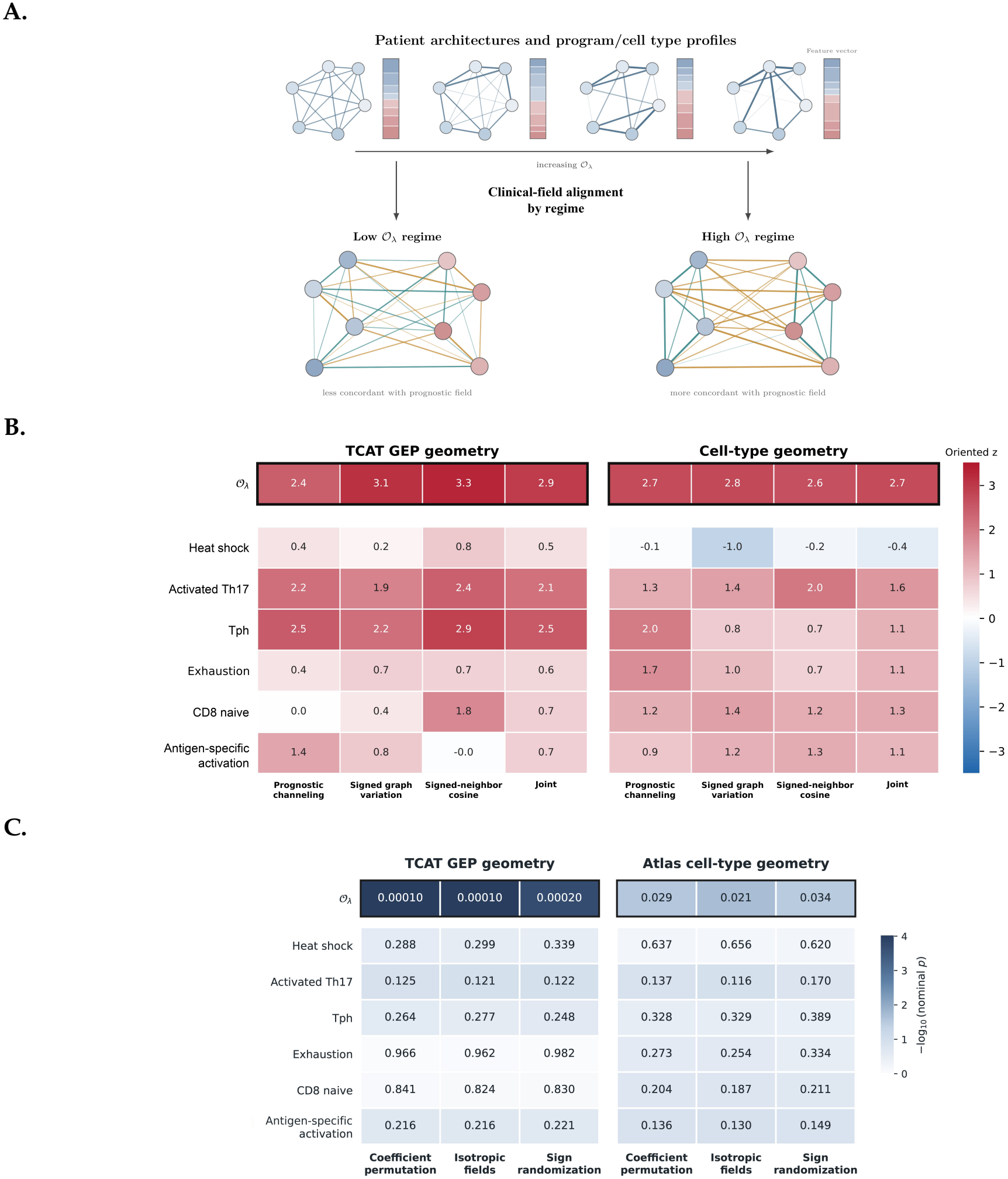
Spectral organization and alignment of the clinical field at two biological resolutions. **A**. Illustrative networks showing gene dependencies within products and feature correlations across products. The clinical field is fixed within each representation, and correlation geometry differs between regimes. **B**. Dataset-adjusted TCAT GEP and atlas phenotype geometries for *O*_*λ*_ and six transition-selected cellular predictors. Contrasts compare the adverse with the favorable regime (low versus high for CD8 naive, high versus low for the other contexts). Positive permutation-standardized scores denote greater channeling, lower variation, or greater neighbor alignment in the adverse regime. The columns report *Z*_*C*_, *−Z*_*V*_, *Z*_*A*_, and their mean *Z*_joint_. Color encodes the oriented, permutation-standardized contrasts. **C**. Clinical-field specificity in the same seven contexts. Cells report nominal one-sided *p* values for the pooled three-metric joint contrast against coefficient-permuted, isotropic, and sign-randomized surrogate fields with the regime correlation matrices held fixed. Color encodes *−* log_10_(*p*) on a shared scale.

Holding ***β*** fixed across spectral regimes, we estimated separate feature correlation matrices from the patients of the high *O*_*λ*_ regime and then the patients of the low *O*_*λ*_ regime, denoted *R*_*g*_ = (*r*_*jk,g*_) for each regime *g*, using probabilistic memberships from the transition survival model. We took the arithmetic mean of the correlation matrices across bootstrap realizations (the arithmetic mean here preserves positive semidefiniteness Methods; Supplementary diagnostics).

To quantify the alignment between the clinical field and the feature correlation geometry, we introduced three statistics. First, *prognostic channeling C*_*g*_ was defined as

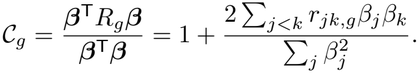

Here, a pair of features increases channeling when the two features are positively correlated and have the same clinical direction, or when they are negatively correlated and have opposing clinical directions. We can also view prognostic channeling as simply the Rayleigh quotient of the regime correlation matrix *R*_*g*_ evaluated at the clinical field ***β*** [33]. Prognostic channeling equals one when features are uncorrelated and increases when their correlations reinforce a common clinical direction.

Second, *signed graph variation* measures agreement of clinical effects across correlated feature pairs. Within regime *g*, let

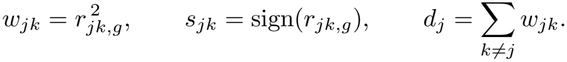

Like prognostic channeling, signed graph variation evaluates a Rayleigh quotient of the clinical field. While prognostic channeling summarizes the combined contribution of all feature correlations to variation along the clinical direction, signed graph variation measures agreement between the clinical coefficients of connected features after accounting for the sign of each edge. We define *V*_*g*_ using the degree-normalized Rayleigh quotient of the signed graph Laplacian [34],

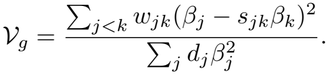

Hence, signed graph variation is lower when correlated features have similar clinical effects and negatively correlated features have clinical effects of opposite direction.

Third, *signed-neighbor cosine* measures agreement between each feature’s clinical effect and the collective orientation of its correlated neighbors. For *d*_*j*_ *>* 0, let

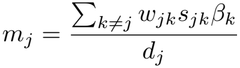

be the clinical effect implied by the signed neighborhood of feature *j*. Define signed-neighbor cosine *A*_*g*_ as

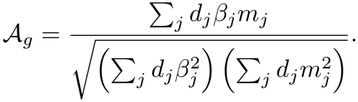

Whereas signed graph variation compares individual pairs, neighbor cosine first combines the contributions of each feature’s neighbors; higher signed-neighbor cosine values indicate that each feature’s clinical direction is more consistent with the combined pattern of its neighbors. And so, these three statistics examine whether clinical alignment happens at the level of 1) overall correlation structure, 2) individual pairs, and 3) their broader neighborhoods.

Although correlation structures were broadly similar between regimes (Pearson correlation between corresponding off-diagonal matrix entries: *r* = 0.914 for TCAT GEPs and *r* = 0.906 for atlas cell types), their alignment with the clinical field differed. We found that the high *O*_*λ*_ regime exhibited greater channeling, lower signed graph variation, and stronger neighbor alignment in both the TCAT GEP and atlas cell type representations (Fig. 5B). The high-minus-low contrasts (Δ*C,ΔV,ΔA* ) were (1.035, − 0.138, 0.122) for TCAT programs and (0.548, − 0.234, 0.170) for atlas cell types. To assess the combined evidence from our three alignment statistics, we standardized each observed high-minus-low contrast against its within-dataset membership-permutation distribution to obtain a *z* score, and then combined the three scores into a single alignment statistic,

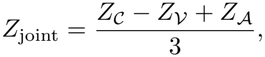

where the minus sign ensures that lower signed graph variation contributes to stronger alignment, alongside greater channeling and higher signed-neighbor cosine. We computed the same average for each permutation to obtain the joint null distribution. The observed joint scores were 2.93 for TCAT GEPs and 2.71 for atlas cell types (two-sided permutation *p* = 0.0008 and 0.0014, respectively). Thus, cellular programs and cell types varied together more coherently along a common clinical risk direction in the high *O*_*λ*_ regime.

We repeated this analysis for the six cellular features whose transition survival models had 95% credible intervals excluding a hazard ratio of one. For GEP-defined contexts, we excluded the defining GEP from its own geometry. For ASA, we excluded its four constituent GEPs. Of the seven contexts tested, spectral organization showed the strongest evidence of a difference in clinical alignment in both representations (Benjamini–Hochberg-adjusted two-sided *p* = 0.0056 for TCAT and *p* = 0.0098 for atlas composition). Tph also showed evidence in GEP geometry (adjusted *p* = 0.0287). Only spectral organization remained significant after adjustment in the joint test for both geometries.

Next, we withheld each dataset in turn. The other seven datasets supplied both the *O*_*λ*_ transition used to assign held-out memberships and the individual continuous-survival coefficients defining the clinical field. We evaluated that field on the held-out cohort’s feature correlations. For the cross validation, we used the five cohorts (in total 144 patients) that had sufficient membership weight in both held out regimes (Supplementary Table S4). In both representations, four of the five held out cohorts showed the expected overall pattern of stronger clinical alignment in the high *O*_*λ*_ regime. The combined one-sided permutation test supported transfer of the alignment contrast to held-out cohorts for atlas composition (*p* = 0.011). The TCAT contrast had the same direction (*p* = 0.103; Supplementary analysis).

We next asked whether the regime contrast was specific to the observed clinical direction. We held the high and low regime correlation matrices fixed and compared the learned field with three families of surrogate fields. *Coefficient permutation* reassigned clinical coefficients among features, *isotropic sampling* generated random directions and scaled them to the norm of the learned field, and *sign randomization* independently flipped the signs of the clinical coefficients. After applying the same transformations for compositional constraints and feature scaling to all fields, we used one-sided tests to assess how often a surrogate contrast matched or exceeded the observed contrast in the direction identified by the primary analysis (Supplementary methods).

We ran all three surrogate tests with *O*_*λ*_ and the six cellular comparator contexts (Fig. 5C). Full cohort *p* values were adjusted across the seven contexts within each representation, statistic, and surrogate family. For the *O*_*λ*_ regime comparison, the TCAT three-metric joint test gave nominal *p* = 0.0001 for coefficient permutation and isotropic fields (adjusted *p* = 0.0007 for each) and nominal *p* = 0.0002 for sign randomization (adjusted *p* = 0.0014). In the atlas cell type geometry, the channeling test gave the strongest evidence, with nominal *p* = 0.0047, 0.0029, and 0.0148 for coefficient permutation, isotropic fields, and sign randomization, respectively (adjusted *p* = 0.033, 0.020, and 0.104). The atlas three-metric joint test gave nominal *p* = 0.0292, 0.0212, and 0.0337 in the same order (adjusted *p* = 0.204, 0.148, and 0.236). Among the seven contexts, *O*_*λ*_ yielded the smallest joint-test *p* value in both geometries under all three surrogate families. Even the strongest joint surrogate result among the six cellular comparators, activated Th17 in atlas geometry under isotropic sampling, only gave nominal *p* = 0.116 (adjusted *p* = 0.304), which was well above the joint *p* values for *O*_*λ*_ in either geometry. Tph was the only cellular comparator to meet the adjusted threshold in the primary joint membership-permutation test, yet its TCAT joint surrogate tests gave little evidence of specificity to the learned clinical direction (nominal *p* = 0.248–0.277, adjusted *p* = 0.404–0.434). Therefore, *O*_*λ*_ stood out among the contexts we tested for how clearly cellular features varied together in alignment with the observed clinical direction in the high *O*_*λ*_ regime.

We then repeated all three surrogate tests in the held-out *O*_*λ*_ comparisons. In each fold, we used the clinical field estimated from the seven training cohorts and held the evaluated cohort’s regime correlation matrices fixed. Across held-out cohorts, coefficient permutation, isotropic fields, and sign randomization gave nominal joint-test *p* values of 0.0276, 0.0297, and 0.0429 for TCAT GEPs and 0.0463, 0.0457, and 0.0784 for atlas cell types, respectively. Prognostic channeling in TCAT GEP geometry provided the clearest held-out evidence of specificity (nominal *p* = 0.0038, 0.0039, and 0.0062, respectively).

Across both TCAT GEP and atlas cell type geometries, the high *O*_*λ*_ regime showed stronger alignment between the way cellular features varied together and their associations with treatment failure. Thus, the findings of our clinical-field analysis link gene-dependency architecture within individual infusion products to how transcriptional programs collectively align with clinical risk across patients. The agreement across these two descriptions of cellular state suggests that spectral organization reflects an emergent property of how the infusion product is organized as a whole.

### Transcriptional pathways associated with high and low spectral organization

Having examined how spectral organization relates to cellular programs, composition, and the clinical-association field, we next asked how *O*_*λ*_ corresponded to established biological pathways through gene set enrichment analysis (GSEA) [35]. Within each infusion product, we generated cell-level Bayesian bootstrap pseudobulk particles over the STRING genes that were shared across all datasets. For each gene, we estimated the partial Spearman correlation between pseudobulk expression and asymptotic *O*_*λ*_, adjusting for dataset and propagating uncertainty in both estimates (Methods). For the GSEA, genes were ranked by the ratio of their posterior mean correlation to its posterior standard deviation. We analyzed both the Hallmark [36] and Reactome [37] gene sets against the complete eligible ranking. Normalized enrichment scores (NES) were used to assess how gene sets are distributed among the ranking of genes associated with spectral organization. No clinical outcomes were used in the gene associations nor the enrichment analysis.

At a within-collection false-discovery rate (FDR) below 0.05, 31 of 50 Hallmark sets and 386 of 1,131 Reactome pathways were significantly enriched. Hallmark results revealed a broad transcriptional contrast (Fig. 6A); lower spectral organization was associated with E2F targets (NES = −4.50), MYC targets V1 (−3.57), and the G2M checkpoint (−2.88), along with oxidative phosphorylation (−1.97) and DNA repair (−1.69). On the other hand, higher spectral organization was associated with interferon-*α* response (NES = 2.13), IL6– JAK–STAT3 signaling (2.12), interferon-*γ* response (2.00), inflammatory response (2.00), and TNF signaling through NF-*κ*B (1.90).

**Figure 6:**
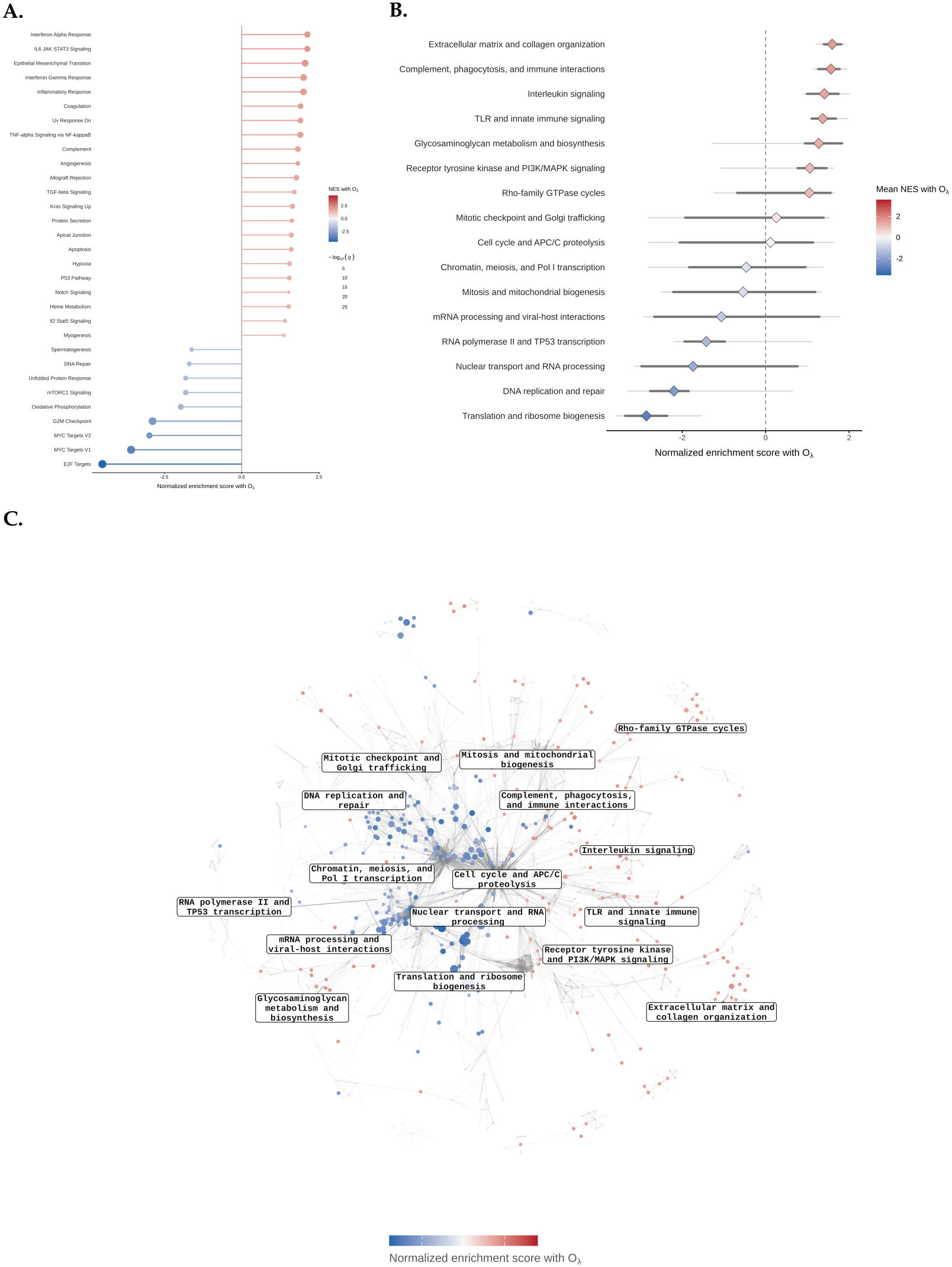
Transcriptional systems associated with spectral organization. **A**. The 31 Hallmark sets enriched at within-collection *q <* 0.05, ranked by normalized enrichment score (NES). Positive NES correspond to higher 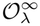 and negative NES to lower values. Color shows NES; point size shows *−* log_10_(*q*), capped at 25. **B**. NES summaries for all pathways within each labeled Reactome community. Diamonds show means, thick segments show the 10th–90th percentile range, and thin segments show the full range of pathway scores. **C**. Reactome network with edges and node positions defined by pathway overlap in gene membership. Color shows NES; node size and opacity reflect within-Reactome enrichment evidence. Communities with at least six pathways meeting *q <* 0.05 are labeled.

To further examine relationships among the Reactome results, we constructed a pathway-overlap graph from all tested pathways and grouped them by shared gene membership [38] (Fig. 6C; Methods). Pathway overlap determined the communities and network layout; enrichment scores were overlaid only after this structure had been defined. The resulting communities thus summarize related pathways with overlapping gene membership.

Lower *O*_*λ*_ was most consistently associated with translation and ribosome biogenesis, DNA replication and repair, nuclear transport and RNA processing, and pathways involved in RNA polymerase-associated transcription (Fig. 6B). All 16 pathways in the translation and ribosome-biogenesis community had negative NES, with mean NES = − 2.86. The 43-pathway DNA replication and repair community was also predominantly negative, with mean NES = − 2.20. Broader communities involving cell-cycle control, mitochondrial biogenesis, and Golgi trafficking contained pathways in both directions. The negative associations were therefore concentrated more specifically within biosynthetic, replicative, and genome maintenance processes.

This gene-level cell-cycle enrichment does not imply a uniform increase in the TCAT proliferation score, which was only weakly related to *O*_*λ*_. Rather, the two analyses capture different aspects of proliferation; TCAT summarizes the relative usage of three predefined cycling programs, whereas enrichment tests whether genes belonging to cell-cycle systems concentrate toward one end of the full gene-association ranking. The two results therefore describe related but nonidentical aspects of proliferative biology.

Higher *O*_*λ*_ was most consistently associated with immune signaling and interactions with the extracellular environment (Fig. 6B). All pathways had positive NES in the communities for interleukin signaling (18 path-ways, mean NES = 1.41), Toll-like receptor and innate immune signaling (20 pathways, 1.37), complement, phagocytosis, and immune interactions (17 pathways, 1.57), and extracellular matrix and collagen organization (19 pathways, 1.60). Pathways related to receptor tyrosine kinase and PI3K–MAPK signaling (54 of 56 pathways, mean NES = 1.06), Rho-family GTPase cycles (19 of 22, 1.05), and glycosaminoglycan metabolism and biosynthesis (24 of 25, 1.27) were also predominated positive in score. Hence, these associations show that more persistent gene-dependency architectures may correspond to cellular systems involved in signal reception and propagation, membrane and cytoskeletal regulation, and interactions with the extracellular environment.

These GSEA results raise the possibility that spectral organization reflects an infusion product’s capacity to reconfigure. However, further investigating this hypothesis would require tracking cellular organization over time or following a defined perturbation. The enrichment results instead provide product-level transcriptional context for this hypothesis and identify pathways whose activity and role in shaping gene-dependency architecture could be examined in such studies. Moreover, these findings do not imply that signaling is uniformly adverse or biosynthesis uniformly favorable; the consequences of these systems likely depend on their cellular context. The complete enrichment results, leading-edge genes, and multiple-testing sensitivities are reported in Supplementary analysis and the accompanying supplementary tables.

## Discussion

The findings of our analysis connect gene-dependency architecture within CAR T infusion products to the collective clinical organization of cellular features across patients. Higher *O*_*λ*_ was associated with increased treatment failure risk and identified a regime in which feature correlations aligned more strongly with a common clinical field. These results address two aspects of the same systems-level question: whether infusion-product organization contains clinically relevant information beyond the abundance of individual cellular states. The connection spans distinct observational scales; spectral organization is inferred from gene dependencies across cells within each patient, whereas clinical-field organization is measured from variation among patients. And so, their correspondence is an empirical finding rather than a direct consequence of the mathematical definition of *O*_*λ*_.

The clinical-field analysis illuminates how this correspondence manifests. Holding individual feature associations with treatment outcome fixed, we compared feature correlations between spectral regimes. Here, we found that in the high *O*_*λ*_ regime, features associated with the same clinical direction varied together more coherently, while features with opposing associations more often varied in opposite directions. Both TCAT programs and atlas composition showed this contrast in the full cohort, which provides the primary evidence of this observation. This primary result was then supported by two secondary sensitivity analyses. First, in the held-out-dataset evaluation, atlas composition showed clear validation evidence, and TCAT programs showed suggestive, aligned statistical evidence. Next, we used compositional surrogate analyses to examine whether the regime contrast was specific to the observed clinical field, where we retained the correlation matrices and evaluated the clinical-field analysis on alternative clinical fields that accounted for compositional constraints and feature scaling. In this analysis, specificity was strongest for TCAT program geometry, while atlas phenotype specificity was supportive yet consistently resolved after multiplicity adjustment and across surrogate constructions. These analyses provide complementary evidence. Atlas composition showed the clearest transfer of the membership contrast across cohorts, while TCAT showed the strongest specificity to the observed clinical direction.

The survival results support spectral organization as a clinically relevant dimension of infusion product biology. Higher *O*_*λ*_ was associated with increased treatment failure hazard both continuously and across spectral regimes, with an estimated twofold difference in hazard between high and low regimes. Furthermore, the survival association with spectral organization persisted across dataset omission and cross-fitted survival evaluation. *O*_*λ*_ bootstrap estimation also retained the general association, with slightly stronger continuous hazard, supporting its robustness to the asymptotic estimation procedure. Finally, its relationship to CAR T treatment outcome remained directionally consistent when modeled jointly with detailed transcriptional programs or cellular composition. The prognostic relevance of *O*_*λ*_ is particularly striking given that it is defined entirely by statistical dependencies among genes. Nonetheless, it is able to capture some underlying principle of immune function and reflect how the infusion product responds as a system, even without predefining any cellular states or immune programs.

The Bayesian survival framework used in our analysis was designed to assess association with treatment outcome while accounting for heterogeneity between cohorts. For a given feature, the model estimated a population-level survival association, along with dataset-specific effects, which allowed us to infer both a feature’s overall clinical direction and the consistency of that feature’s clinical relevance across cohorts. The estimated associations thus account for measured differences in disease and CAR T product across the included cohorts. Thus, a further contribution of this work is the survival framework itself, independent of its application to spectral organization.

The relevance of *O*_*λ*_ association with treatment outcome further raises the question of how a patient’s transcriptional network organization should be interpreted biologically. Again, the networks here are computed from statistical dependencies and thereby do not establish molecular interactions, regulatory connections, or communication between cells. Hence, interpreting *O*_*λ*_ in terms of both known and emergent biology requires careful consideration. To start, spectral organization was related to, but incompletely accounted for by, the measured biological components of the product. That is, the correlation associations between *O*_*λ*_ and the individual programs and phenotypes were generally modest, and *O*_*λ*_ was not convincingly recovered by the multivariate reconstruction, nor by composition-matched synthetic products. Similarly, technical covariates provided an incomplete account of *O*_*λ*_. These observations motivate considering spectral organization as a candidate emergent property of the infusion product, as it is collective by construction and summarizes dependency patterns across the entire gene network.

The spectral benchmarks help clarify what our introduced *O*_*λ*_ contributes beyond established summary statistics from the field of spectral graph theory. The two other summary statistics we considered were the spectral gap, which only characterizes the slowest nonstationary mode of a graph’s spectrum, and the normalized Kemeny’s constant, which gives equal weight to the relative lifetimes of all nonstationary modes. The weighting we introduce instead gives the greatest influence to modes whose lifetimes depart most from the equal-weight complete-graph reference, which we hypothesized carried the bulk of biologically relevant signal. A higher value of *O*_*λ*_ therefore indicates that the modes that most distinguish the gene-dependency network from the reference tend to persist longer under graph diffusion. When we compared all three summaries using the same patient-bootstrap analysis, the association with treatment failure was clearest for *O*_*λ*_ (Supplementary Fig. S2). This suggests that in the contexts of these CAR T infusion products, *O*_*λ*_ appears to illuminate some biologically relevant behavior that the others do not quite capture.

The transcriptional associations from the GSEA do provide meaningful biological context in through which this organization occurs. Higher spectral organization was accompanied by increased signatures of signaling, inflammatory, and environmental-interaction systems, whereas lower values accompanied translation, replication, RNA processing, and genome maintenance. However, these product-level expression associations do not directly measure pathway activity or identify mechanisms responsible for the dependency network. For example, we found both cell-cycle gene set enrichment associated with lower *O*_*λ*_ despite a negligible relationship between spectral organization and the TCAT proliferation score, which illustrates how gene-level enrichment and relative program usage can capture different aspects of the product. The consequences of signaling and biosynthesis likely also depend on the specific processes involved and their cellular context.

One hypothesis is that spectral organization retains an imprint of the product’s history that also relates to its capacity for future reconfiguration. Disease and treatment exposures may shape the cellular states entering apheresis [39], while stimulation and expansion during manufacturing can alter their differentiation and functional potential [40]. These processes may also preserve or reorganize dependencies among cellular programs. The architecture observed at infusion could therefore retain information about earlier conditions while identifying the configurations from which subsequent responses develop. To that end, historical imprint and future plasticity are compatible interpretations, which positions spectral organization as a potential link between the product’s history and its capacity for subsequent reconfiguration. One possibility is that the dependency patterns captured in high *O*_*λ*_ products constrain which cellular programs can change independently after infusion. Across patients in the high *O*_*λ*_ regime, programs with adverse clinical associations tended to vary together. Thus, if those programs also remain coupled as cells respond to stimulation, a product that starts in an unfavorable configuration may have more difficulty shifting toward a state that supports an effective response. Future work could combine longitudinal profiling with controlled perturbations to assess when these dependencies emerge, how they persist, and whether they anticipate changes in cellular state and function.

The myeloid-associated findings provide a plausible biological basis for this hypothesis. Deng et al. identified rare monocyte-like cells in manufactured CAR T infusion products that were associated with severe neurotoxicity; however, their cellular lineage remained unresolved [11]. Similar myeloid-like populations were subsequently detected in an independent study of axi-cel infusion products [41]. Before manufacturing, Carniti et al. traced an outcome-associated myeloid signature to monocyte–T-cell complexes in leukapheresis products [42]. Experimental manufacturing work further showed that high monocyte content can impair CAR T-cell production and that monocyte depletion can improve transduction and expansion [43]. Perhaps most notably, evidence suggests that mixed-lineage transcriptional features can arise through biological responses to stimulation and cellular interaction. In an infection setting, Kang et al. found mixed T-cell and monocyte transcriptional signatures in individual cells that had been physically separated from complexes, which suggests that T cells can acquire exogenous transcriptional signals from cell-to-cell interactions [44]. Similarly, Li et al. identified CAR T-cell populations with non-T-cell transcriptional features after infusion and showed experimentally that excessive antigen stimulation or suppressed TCR signaling could induce B-cell-associated features [45]. Therefore, these prior studies provide evidence that T cells can adopt mixed-lineage transcriptional patterns, and these mixed-lineage signals are indeed biologically relevant. The association of myeloid-like signals with higher *O*_*λ*_ raises the possibility that traces of earlier cellular experiences are reflected in which gene dependencies are detected across infusion cells, leaving an imprint on the product’s transcriptional architecture.

It is also worth noting mixed-lineage features did not generally share a uniform association with spectral organization in our analysis. The stronger myeloid-associated signal may therefore reflect a particular cellular context rather than a general effect of mixed-lineage expression. Consequently, if these signals reflect a product-wide transcriptional configuration marked by competing programs rather than coordinated ones, their presence may constrain which cellular states the product can adopt. In this view, myeloid-like signals are one possible manifestation of the broader transcriptional organizational schemes that *O*_*λ*_ measures. The removal of predicted doublets during preprocessing, together with the clinical and experimental evidence for mixed-lineage features, thereby supports a biological interpretation of the retained signals.

The systems-level contribution of our framework does not depend on resolving whether or not the prognostic and organizing qualities of *O*_*λ*_ are truly emergent beyond what the individual components of the cellular system can explain. Spectral organization nevertheless provides a common context in which established cellular programs and phenotypes can be examined together in relation to clinical outcome. Its relevance extends beyond its own survival association; it is a profound observation that a property inferred from gene dependencies, independent of any prior knowledge of biological gene/program associations, can distinguish how cellular features collectively align with clinical risk across patients. Even if spectral organization can ultimately be explained by the complete description of cellular states or programs, it still provides an informative summary of how those states are organized collectively. As it stands now, our framework can be interpreted as an organizing framework for CAR T infusion product biology, while its status as an emergent biological property remains open.

Several inferential boundaries of our framework do still remain. First, the cohorts were retrospective, and independent estimates from the smallest datasets remained uncertain. Furthermore, the inferred spectral asymptote is an extrapolated limit under a graph-reconstruction procedure where delayed trajectory changes can be difficult to resolve, and the estimated uncertainty is conditional on the selected model. The spectral regimes are also probabilistic, outcome-associated contexts rather than established biological thresholds. Finally, the clinical field comprises associations from individual feature models, not causal effects or a jointly validated patient-level predictor.

Generalization to other biological systems also requires attention to their cellular composition. CAR T infusion products provide a comparatively restricted cellular system, composed largely of related T-cell states. In tissues/contexts containing more divergent cell types, differences in baseline gene expression and detection frequencies can generate strong dependencies when cells are pooled, potentially dominating the network spectrum. The present construction may likely require adaptation, such as estimating dependencies within lineages or accounting for differences between cell types, before its interpretation can be extended to other systems. Although we hypothesize that collective organization and reconfigurability are relevant beyond CAR T-cell therapy, their measurement will depend on the biological system and the scale at which it is represented.

It is also worth mentioning that our framework characterizes the infusion product without directly profiling the patient’s tumor or its microenvironment. Even though the survival models accounted for disease and treatment, they did not incorporate patient tumor burdens or molecular subtypes. The observed associations therefore indicate that the infusion product itself carries information relevant to treatment outcome that remains across clinical contexts. This does not imply independence from tumor biology; the product may retain an imprint of prior disease exposure, and its subsequent behavior may depend on the environment it encounters. An interesting future direction would be pairing infusion product architecture with measurements of tumor state and the surrounding tumor microenvironment. Such analyses could examine whether a given dependency architecture supports effective responses in some tumor contexts but encounters constraints in others, connecting product-level organization to the reciprocal interactions through which treatment response develops.

CAR T infusion products contain clinically relevant information in both their constituent states and the dependency structures in which those states coexist. Spectral organization connects these levels by relating within-product gene dependencies to the collective clinical organization of programs and cell types across patients. This connection provides a basis for studying familiar biological features as parts of clinically meaningful configurations. It also motivates investigating how a product’s history shapes its architecture at infusion and whether that architecture helps explain its subsequent response to treatment.

## Conclusion

Spectral organization identifies a clinically relevant dimension of CAR T infusion products: the dependency architecture in which their cellular features coexist. Higher *O*_*λ*_ was associated with increased treatment failure risk and with stronger alignment of cellular feature correlations along a common clinical direction. This empirical connection links statistical gene dependencies within individual products to the collective organization of transcriptional programs and cellular composition across patients. Established cellular features retain their biological importance, while spectral organization reveals a context in which their clinical associations become collectively expressed. These findings support studying CAR T products as complex cellular systems whose architectures may carry an imprint of prior conditions and bear on subsequent responses. Understanding their therapeutic potential therefore calls for examining both which cellular states are present and how they are organized together. Our framework is a first step toward understanding that organization.

## Methods

### Cohort inclusion and clinical outcome harmonization

We included publicly available datasets containing single-cell RNA-sequencing profiles of CAR T infusion products and patient-level clinical outcome information. Patients were retained in the analysis when they had both treatment outcome annotations and at least 1,000 retained infusion product cells after preprocessing. The patients spanned diseases of lymphoma and pediatric B-cell acute lymphoblastic leukemia, along with CAR T product types including axi-cel, tisa-cel, and tandem CD19/CD20 products. The final analysis population included 163 patients across eight cohorts (Table 1).

We harmonized response annotations and follow up information across studies to define treatment failure as primary nonresponse, progression, or relapse. Partial response counted as nonresponse in all cohorts except Maurer, where it counted as response (based on the paper’s specification of categorizing partial response in with responders). Patients who responded and had no subsequently documented event were right-censored at their last available event-free assessment. For any patient with infusion retreatments (Haradhvala patient 29), only the initial treatment and its clinical outcome were analyzed, and any subsequent infusion and outcome were excluded.

All times were measured in months from CAR T infusion. Interval-censored failures were recorded between the last documented event-free assessment, and the first assessment documenting treatment failure. If only one assessment was available, failure was assigned to the interval from infusion to that assessment, and patients without failure were right-censored at that time. For right-censored records, we set the event lower bound to zero by convention and the upper bound to the censoring time.

For continuously reported event times, we obtained month-level intervals by rounding down to the nearest whole month for the lower bound and up to the nearest whole month for the upper bound. The survival likelihood distinguished interval-censored events, exactly observed events, and right-censored observations (Bayesian survival analysis). Models included product and disease covariates, dataset-specific baseline hazards, and partially pooled dataset-specific predictor effects.

### Single-cell preprocessing and quality control

Only infusion-product cells were included, regardless of CAR expression or lineage annotation. For the Bai dataset containing both basal and stimulated conditions, we included only basal cells. Count matrices were obtained from the designated count representation in each source dataset. Gene identifiers were stripped of surrounding whitespace, and exact duplicate gene names were collapsed by summing their counts. Duplicate cell identifiers were made unique where necessary. Cells were retained when at least 200 genes were detected and mitochondrial transcripts comprised less than 10% of total counts. Mitochondrial genes were identified by the `MT-` prefix.

Scrublet [1] was run independently within the specified sample batches, using patient identity as the default batch definition. Cells classified as predicted doublets were removed. The 1,000-cell patient inclusion threshold was applied to the retained infusion product cells after preprocessing. All preprocessing was done with Scanpy [2].

### STRING gene harmonization

Gene features were harmonized to preferred human gene symbols using the STRING v12.0 protein-information and protein-alias files for taxon 9606 [3]. Preferred symbols were matched exactly or without regard to case. When no preferred-symbol match was available, an alias was accepted only if it mapped to a single preferred STRING symbol. Unmapped features and aliases mapping to multiple symbols were excluded.

When multiple source features mapped to the same STRING symbol, preferred symbol matches were prioritized, followed by a unique punctuation equivalent alias when available. Among the remaining eligible features, a sole nonzero expression column was retained, and identical nonzero columns were represented once. We summed expression columns for three curated alias groups: SEPTIN4 (SEPT4 and C17ORF47), RAMAC (FAM103A1 and RAMMET), and MTLN (LINC00116 and SMIM37). Each sum included the preferred symbol’s column when present.

### Cell subsampling and construction of gene mutual information graphs

Gene mutual-information networks were constructed separately for each patient’s infusion product using binary gene detection. Let *Y*_*cg*_ = 1 when gene *g* had a positive count in cell *c*, and *Y*_*cg*_ = 0 otherwise.

For a patient with *N* available cells, we selected 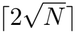 non-full sampling depths and also analyzed the complete sample. We also included fixed anchors at 50, 100, 150, and 200 cells. Additional depths were selected by repeatedly bisecting the largest remaining gaps, with randomized tie-breaking.

Gene eligibility was reassessed within each subsample. Among the STRING-harmonized genes, a given gene was retained when both its detected and undetected states occurred in at least

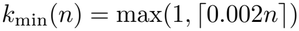

cells. For each pair of eligible genes *g* and *h*, empirical joint and marginal detection probabilities were calculated from the 2 × 2 detection table. Plug-in mutual information was calculated in nats as

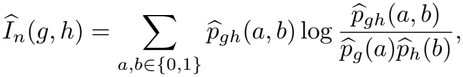

with zero-probability terms contributing zero. At depth *n*, we set 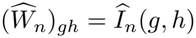 for *g* =/ *h* and 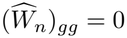. We omit the hat and depth index when describing the graph at a fixed depth. Numerical values were restricted to the valid binary-MI range [0, log 2], and the matrix was symmetrized before spectral analysis. Finite-cell dependence was addressed by modeling the resulting spectral trajectory.

### Spectral organization calculation

Vertices with weighted degree at most 10^*−*12^ were removed before normalization. For the remaining *G* vertices, we formed

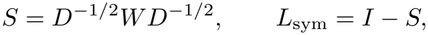

and computed the complete symmetric eigenspectrum of *S*. The largest eigenvalue was removed as the Perron mode.

For a connected graph, the remaining normalized Laplacian eigenvalues are *ν*_*i*_ = 1 − *λ*_*i*_ *>* 0. Their integrated diffusion lifetimes are given by

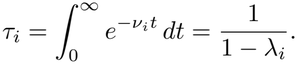

Because *S* has zero diagonal and *λ*_1_ = 1, its nonstationary eigenvalues have mean

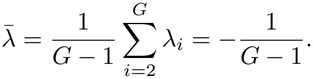

The corresponding reference lifetime is *τ*_ref_ = (*G* − 1)*/G*. Relative lifetimes were therefore defined as

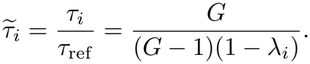

Spectral organization was calculated as

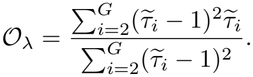

For numerical stability, 1 − *λ*_*i*_ was bounded below by 10^*−*7^ when calculating relative lifetimes.

### Empirical-Bayes inference of asymptotic spectral organization

We transformed spectral organization to its log excess above one:

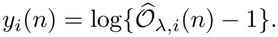

We then modeled each patient’s trajectory using

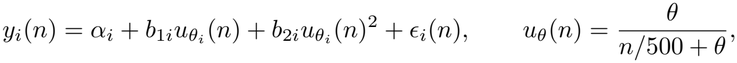

following the asymptotic motivation in the Supplementary Methods. Here *θ*_*i*_ is patient *i*’s convergence timescale. The factor of 500 expresses sampling depth in units of 500 cells, so *u*_*θ*_(*n*) = 1*/*2 at *n* = 500*θ*. The prior median *θ* = 2 therefore corresponds to a characteristic depth of 1,000 cells. Because *u*_*θ*_(*n*) → 0 as *n* → ∞, the corresponding asymptote is

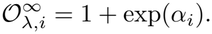

Here, the asymptote describes the extrapolated limit of the graph-reconstruction procedure.

We approximated the positive convergence-timescale distribution with nine equal-probability quadrature nodes under

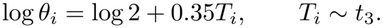

For each node, we used `nlme` [4] to fit a linear mixed-effects model by maximum likelihood. Population-level intercept, slope, and curvature terms were accompanied by correlated patient-specific random coefficients. Residual standard deviation varied as a power function of *n/*500. Of the 20,992 trajectory observations from the 163 patients, the population fits used 4,890 points selected at approximately uniform depths (target 30 per patient). Across components, the asymptote mean and variance were anchored to those fitted at *θ* = 2; conditional slope structure remained component-specific. The trajectory model included no dataset-specific fixed effects or clinical outcomes.

Patient-specific inference used a subset targeting 40 uniformly spaced depths and a Student-*t* residual likelihood with four degrees of freedom. The squared residual scale was the sum of a fitted depth-dependent variance and a depth-independent variance term. The intercept retained a Gaussian population prior, while the two slope coefficients conditional on the intercept followed a bivariate Student-*t* prior with ten degrees of freedom.

To limit information accumulation from overlapping nested subsamples, we tempered each patient’s summed log likelihood by

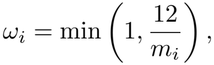

where *m*_*i*_ is the number of retained fitting points. Tempering here limited the contribution of densely sampled trajectories without requiring an estimate of the covariance between depths.

We numerically optimized patient coefficients and approximated their local covariance using the inverse Hessian. Laplace approximations to component marginal likelihoods, combined with equal prior quadrature weights, determined the timescale weights. We held the fitted population parameters fixed during patient-specific inference.

### Posterior uncertainty and particle sampling for each asymptote

For patient *i* and timescale component *q*, we drew the log-excess asymptote as

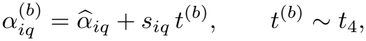

Where 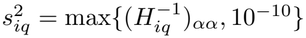 and *H*_*iq*_ is the regularized local Hessian of the negative log posterior at the fitted coefficients. The numerical lower bound protects the covariance calculation. The component scale came directly from this curvature, without a post-hoc calibration floor on asymptote uncertainty. The depth-dependent residual scale described above entered the trajectory likelihood. We generated 50,000 mixture draws per patient using the patient-specific timescale weights and transformed each draw as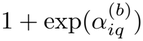.

The reported central estimate was the weighted average of the transformed component estimates,

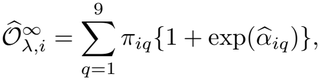

where *π*_*iq*_ denotes the component weight.

The mixture distribution was summarized by its median and 2.5th and 97.5th percentiles. For downstream uncertainty integration, *M* equal-weight particles were selected at midpoint quantiles

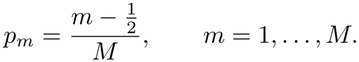

The primary analyses used *M* = 16. For validation against held-out observations at greater cell depths and full-trajectory estimates, see the Supplementary Information.

### Estimating TCAT program usages and scores for each patient

TCAT was applied separately to each dataset using the TCAT.V1 reference implemented in starCAT [5]. For each STRING-harmonized input matrix, inference used the available genes overlapping the TCAT reference. The nonnegative usages of the reference gene expression programs were estimated in each analyzed single cell. The analyzed TCAT features comprised 52 GEPs and two derived functional scores; TCAT categorical cell type annotations were not used. All GEP usages sum to one within each cell.

For patient *i*, let *u*_*icj*_ denote the usage of program *j* in cell *c*. The central patient-level program composition was

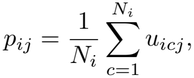

where *N*_*i*_ is the number of retained infusion cells. We transformed the composition **p**_*i*_ = (*p*_*i*1_, …, *p*_*iK*_) using

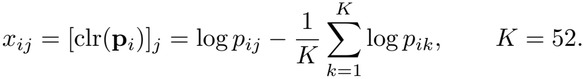

Each transformed usage therefore represents enrichment relative to the geometric mean of the patient’s GEP composition. The transformation was applied after patient-level aggregation, rather than to individual cells.

Antigen-specific activation (ASA) and proliferation scores were derived from the transformed GEPs rather than from the original cell-level composite scores. ASA was the mean CLR usage of the TIMD4/TIM3, ICOS/CD38, CTLA4/CD38, and OX40/EBI3 programs. Proliferation was the mean CLR usage of the CellCycle-G2M, CellCycle-S, and CellCycle-Late-S programs. These scores quantify the average relative enrichment of their constituent programs and were not treated as additional coordinates of the GEP composition.

Finite-cell sampling uncertainty was represented by Bayesian bootstrap particles. For each patient and particle *m* = 1, …, *M*, we drew

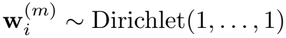

over the patient’s *N*_*i*_ cells and calculated

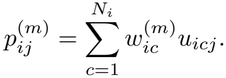

Each weighted composition was CLR-transformed, and ASA and proliferation were recalculated from the corresponding transformed components. The same cell weights were used for every GEP within a particle, preserving dependence among programs and derived scores. We used *M* = 16 equally weighted particles per patient for the downstream uncertainty-aware analyses.

### Atlas cell type annotation and patient-level proportions

Cellular phenotypes were assigned using an atlas-derived scANVI model [6] adapted from the Functional CAR-T Atlas workflow [7]. We used the released V4 reference (`Python_scVI_adata_big_V4_state4.h5ad`), with `manual_celltype_annotation_high` providing the reference phenotype labels and `Product_norm` defining reference batches. The reference contained 11 phenotypes: apoptotic T cells; CD4 central-memory, cytotoxic, and effector memory cells; CD8 cytotoxic, effector memory, and memory cells; monocyte-like T cells; proliferative T cells; regulatory T cells; and ribosomal-enriched cells.

To obtain a common model feature panel, we first restricted the reference to genes present in every processed dataset in the analysis. Genes detected in fewer than 10 reference cells were removed. Within the remaining shared gene universe, we used Scanpy to select 2,000 highly variable genes from the reference using the Seurat v3 method [8]. We ran `seurat_v3` with `Product_norm` as the batch variable and a smoothing span of 0.6. Thus, variability-based feature selection used the reference data, while gene availability across query datasets determined the eligible feature universe.

Reference training followed the atlas authors’ scVI-to-scANVI workflow using scvi-tools version 1.4.2. Raw counts were modeled with a negative-binomial likelihood. The scVI architecture used two layers, encoded covariates, layer normalization in the encoder and decoder, and no batch normalization. The scVI model was trained using the software’s data-dependent default epoch limit, converted to scANVI, and trained for a further 20 epochs using the atlas labels. This procedure produced the atlas-derived classifier used in our analysis.

Each query dataset was mapped separately through scANVI/scArches query adaptation. Raw counts were taken from the counts layer when available, or otherwise from the input expression matrix. Query batches were defined by patient identifiers prefixed with the dataset identity and a query-specific designation, ensuring that they were treated as new batches. Query adaptation used 100 epochs, frozen dropout, zero weight decay, and a batch size of 256. Reference training also used a batch size of 256. The random seed was set to zero. Clinical outcomes were not used for model training or phenotype assignment. For each cell, the annotated object retained the predicted probability vector over the 11 biological phenotypes and the label with the largest probability.

Let *z*_*ic*_ denote the assigned phenotype of cell *c* from patient *i*, and let *N*_*iℓ*_ be the number of cells assigned to phenotype *ℓ*. For the *L* = 11 phenotypes, central patient-level proportions were calculated with a symmetric half-count correction:

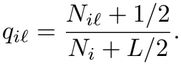

These proportions were CLR-transformed within the atlas composition, separately from the TCAT GEP composition.

Atlas composition uncertainty was represented by Bayesian-bootstrap particles using the same construction as for categorical patient-level features. For each patient, we drew cell weights 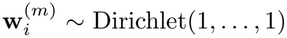 and calculated

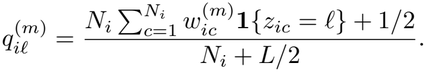

The factor *N*_*i*_ places the weighted proportions on the cell-count scale before adding the half-count correction.

Each particle was then transformed as

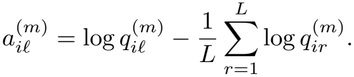

A single cell-weight vector was shared across all phenotypes within each particle, preserving compositional dependence. We used *M* = 16 equally weighted particles per patient. The primary survival analyses retained patients with at least 1,000 infusion cells and marginalized the survival likelihood over these particles. The particles capture uncertainty from sampling a finite number of cells. The classifier, phenotype definitions, and cell labels were held fixed across draws.

### Hierarchical Bayesian survival model for continuous predictors

Outcome associations were estimated using hierarchical Bayesian piecewise-exponential survival models. Follow up was divided at 0, 1, 3, 6, 9, 12, 18, 24, 36, 60, and 120 months, which were chosen based on the observed patient event/censoring timescales. Each individual continuous model included one biological predictor, together with clinical adjustment covariates.

### Predictor standardization and uncertainty particles

For each predictor, the mean 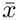 and sample standard deviation *s*_*x*_ were calculated from the central patient estimates across the complete analysis cohort. All predictor particles were transformed using the same scaling,

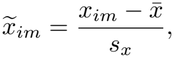

where *x*_*im*_ denotes particle *m* for patient *i*. Predictors were standardized globally rather than within datasets. Patient-level predictor uncertainty was represented by *M* particles with weights *w*_*im*_ satisfying

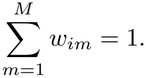

For spectral organization, particles represented the inferred distribution of *O*_*λ*_ from the trajectory model. For TCAT features, particles represented finite-cell sampling uncertainty estimated by the patient-level Bayesian bootstrap. For atlas cell type features, each cell retained its assigned atlas cell type across bootstrap draws.

Within each patient, we varied the weights assigned to individual cells and recalculated the resulting cell type proportions in each draw. The particles for *O*_*λ*_, TCAT features, and atlas cell type proportions were generated before fitting the survival models and were used as model inputs. These first stage distributions were constructed before survival model fitting. We used *M* = 16 in the primary analyses, chosen based on previous numerical comparisons with higher particle resolutions.

### Continuous predictor effects and clinical covariates

For patient *i* from dataset *d* during time interval *k*, the conditional hazard was modeled as

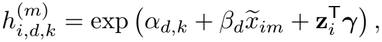

where *α*_*d,k*_ denotes the dataset-specific baseline log hazard, *β*_*d*_ is the dataset-specific predictor coefficient, and **z**_*i*_ contains the treatment and disease indicators. Treatment categories included axi-cel, tisa-cel, and tandem CD19/CD20 CAR T products; disease categories included lymphoma and pediatric leukemia.

### Hierarchical priors

Predictor coefficients were partially pooled across datasets:

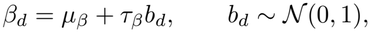

with priors

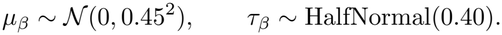

Clinical adjustment coefficients had independent priors

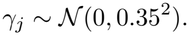

Each dataset received an independent initial baseline log hazard,

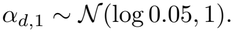

Subsequent baseline log hazards followed a Gaussian random walk:

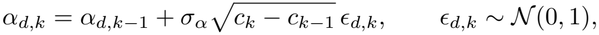

where *c*_*k*_ is the midpoint of time interval *k* in months and

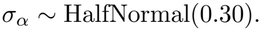

The smoothness parameter was shared across datasets, while each dataset had its own initial log hazard and changes in log hazard between successive intervals. Predictor and clinical adjustment coefficients were constant over follow up, giving a proportional-hazards model conditional on dataset and predictor value.

### Survival likelihood and uncertainty integration

For patient *i* with predictor value *x*, let *h*_*i*_(*t*| *x*) denote the hazard, *H*_*i*_(*t*| *x*) the cumulative hazard, and *S*_*i*_(*t* |*x*) = exp {−*H*_*i*_(*t* |*x*)} the survival function. Let *a*_*i*_ and *b*_*i*_ denote the lower and upper bounds of the reported event/censor interval. An event known to occur in (*a*_*i*_, *b*_*i*_] contributed

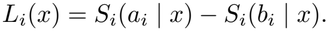

A patient who remained event-free through the censoring time *b*_*i*_ contributed

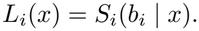

An event observed exactly at time *t*_*i*_ contributed

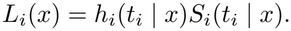

For an interval-censored event, the likelihood is the probability that the event occurred anywhere within the reported interval.

Predictor uncertainty was incorporated by marginalizing each patient’s likelihood over the supplied particles:

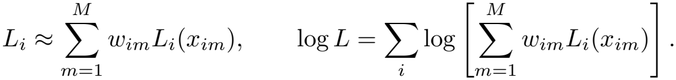

The weighted likelihood sum was evaluated using a numerically stable log-sum-exp calculation. We averaged each patient’s likelihood over their predictor particles.

### Hazard ratio summaries, cohort agreement, and sampling diagnostics

For the continuous survival model, the primary estimand was the population-level hazard ratio exp(*µ*_*β*_) per one global standard deviation increase in the predictor. We reported its posterior median, equal-tailed 95% credible interval, and posterior probability *P* (*β >* 0) = *P* (*µ*_*β*_ *>* 0 | *D*), where *D* denotes the combined clinical and predictor information available across cohorts. Dataset-specific hazard ratios and the between-dataset effect standard deviation were also summarized.

Cross-dataset directional agreement was estimated from joint posterior draws as

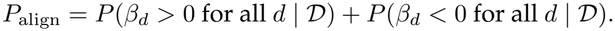

This is the posterior probability that all dataset-specific effects in the hierarchical model share the same sign.

Models were fitted in PyMC [9] using the No-U-Turn Sampler [10] with four chains, 2,000 tuning iterations and 2,000 retained posterior draws per chain, a target acceptance probability of 0.99, and random seed 0.

Sampling diagnostics included divergent transitions, 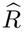, and bulk and tail effective sample sizes [11].

### Bayesian smooth transition survival model

To evaluate regime-based associations with outcome, we replaced the linear feature term in the continuous survival model with a logistic transition. Features included *O*_*λ*_, CLR-transformed TCAT GEP usages and atlas phenotype proportions, and TCAT composite scores on their defined analysis scales.

For patient *i*, let *x*_*im*_ denote particle *m* of the feature distribution, and let *s*_*x*_ denote the sample standard deviation of its central patient estimates across the complete analysis cohort. The transition function used was

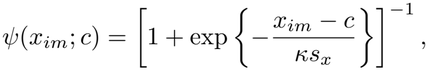

where *c* is the feature-specific crossover location and *κ* = 0.10 is a fixed transition scale in units of the global feature standard deviation. At the crossover, *ψ*(*c*; *c*) = 0.5. We centered the transition at this midpoint in the log-hazard model, so the feature contributed 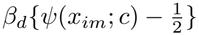 and its contribution was zero at *x*_*im*_ = *c*. For each feature, the crossover was shared across datasets and assigned the prior

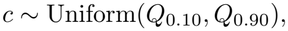

where *Q*_0.10_ and *Q*_0.90_ were calculated from that feature’s central patient estimates in the analysis cohort.

### Smooth transition hazard model and priors

For patient *i* from dataset *d* during time interval *k*, the particle-conditional hazard was

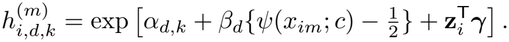

Dataset-specific baseline log hazards *α*_*d,k*_, treatment and disease covariates **z**_*i*_, and their priors were identical to those used in the continuous model. At *x* = *c*, the feature contributes zero to the log hazard. The two limiting regimes contribute − *β*_*d*_*/*2 and +*β*_*d*_*/*2, so reversing their labels reverses the coefficient while preserving the midpoint baseline and its prior. Regime effects were partially pooled according to

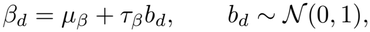

with

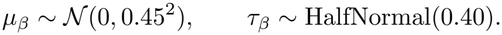

Clinical adjustment coefficients retained independent *N* (0, 0.35^2^) priors. The model permitted either an adverse or protective association with increasing feature values.

The model allowed a smooth transition across predictor values while preserving proportional hazards over time within each dataset. The primary estimand,

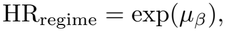

represented the population-level hazard ratio between the limiting high and low regimes, *ψ* = 1 and *ψ* = 0, conditional on the clinical covariates.

### Uncertainty integration and regime membership

Exact events, interval-censored events, and right-censored observations contributed the same conditional likelihoods as in the continuous analysis. For each patient, these likelihoods were marginalized over the predictor particles:

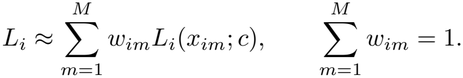

Each particle entered a single smooth hazard function; the model was not a mixture of separate low and high regime survival distributions.

For each posterior crossover draw, soft high regime membership was calculated as

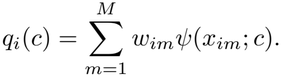

Posterior means and medians of *q*_*i*_(*c*) summarized patient membership while integrating predictor uncertainty and uncertainty in the crossover. The fixed high-regime weight in the clinical-field analyses was *q*_*i*_ = E[*q*_*i*_(*c*) | *D*], calculated by averaging across posterior crossover draws.

### Regime hazard ratios, crossover estimates, and sampling diagnostics

We reported the posterior median and equal-tailed 95% credible interval of the regime hazard ratio and crossover location, together with *P* (*µ*_*β*_ *>* 0 | *D*). Directional agreement across datasets was summarized as

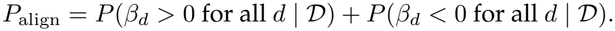

Models were fitted in PyMC using four chains, 2,000 tuning iterations and 2,000 retained posterior draws per chain, a target acceptance probability of 0.99, and random seed 0. Convergence was assessed using divergent transitions, 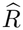, and bulk and tail effective sample sizes.

### Joint predictor models and separate cohort fits

#### Joint models

We evaluated *O*_*λ*_ jointly with either all 52 TCAT GEPs or all 11 atlas phenotype proportions, using both the continuous and smooth transition models. These models retained dataset-specific baseline hazards, partially pooled predictor effects, treatment and disease adjustment, and integration over patient-level feature particles.

In the continuous models, CLR features were centered across patients and divided by a common scale within each composition, defined as the square root of the mean feature variance, which preserved the CLR zero-sum constraint. Population coefficients and dataset-specific deviations were each constrained to sum to zero within the composition. *O*_*λ*_ was standardized separately by its population standard deviation and was not subject to this constraint.

Joint coefficients were regularized using shared shrinkage scales. Before the compositional centering described above, population coefficients and dataset deviations were independent zero-centered Gaussian variables with standard deviations *σ*_global_ and *σ*_dataset_, respectively, with

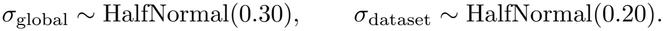

In the smooth transition models, each feature’s crossover was fixed at its individual model posterior median, with transition width equal to 0.10 of its global standard deviation. The joint model included the centered transition values 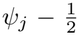 for all predictors, with the same shrinkage priors on their coefficients and no zero-sum constraint. Joint credible intervals were conditional on these fixed crossovers.

We kept cellular features from the same particle draw together to preserve their dependence. Because spectral and cellular particles were generated separately, we paired them at random within each patient. This pairing treated their measurement errors as independent.

Joint hazard ratios describe conditional associations under regularization. Differences from the individual-model HRs can reflect shared information among predictors, coefficient shrinkage, and, for continuous compositional features, differences in scaling.

#### Independent cohort models

To examine cohort-specific directions without partial pooling of survival coefficients, we fitted each predictor separately within each dataset. We evaluated *O*_*λ*_ and cellular features whose population-level 95% credible interval excluded a hazard ratio of one in either the continuous or smooth transition model. Each fit used the same survival likelihood and particle integration as the primary analysis, with a separately estimated baseline hazard and a log-hazard coefficient prior *β*∼ *N* (0, 0.45^2^). Treatment and disease covariates were included only when they varied within the dataset, with coefficient priors N (0, 0.35^2^).

Independent cellular feature models were continuous and used full cohort standardization. For *O*_*λ*_, both the continuous and transition fits used predictor definitions learned from the other seven datasets. For each evaluated cohort, we fixed the continuous center and scale, the posterior-median crossover, and the width (0.10 training SD) before estimating its survival coefficient from that cohort alone. The independent spectral fits used two chains with 1,000 tuning iterations and 1,000 draws per chain (target acceptance 0.97). Leave-one-dataset-out robustness and cross-fitted survival validation are reported separately in the Supplementary analysis. For the independent fits, we summarized direction using each cohort’s posterior-median HR and uncertainty using its 95% credible interval.

### Dataset-adjusted cellular-feature correlations

We estimated correlations among cellular features and between each feature and *O*_*λ*_ using CLR-transformed TCAT GEP usages and atlas phenotype proportions, together with the TCAT composite scores. These analyses used continuous feature values without spectral regime weighting or clinical outcome information.

We used 2,000 draws for TCAT GEP pairs and 4,000 for atlas pairs, atlas–GEP pairs, and associations with *O*_*λ*_. In each draw, we sampled one joint feature particle per patient within each representation. For associations with spectral organization, we independently sampled one asymptote particle per patient. In both analyses, we drew patient weights

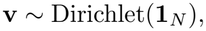

where *N* is the number of patients. Within each draw, we removed dataset-specific weighted means from every feature and *O*_*λ*_:

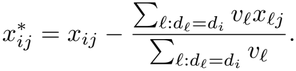

Weighted Pearson correlations were then calculated between the residualized profiles using the same patient weights. This procedure removed between-dataset mean differences while retaining within-dataset variance scales. Treatment and disease were not separately adjusted in these correlation analyses.

For each pair, we reported the median and equal-tailed 95% interval across draws, together with the probabilities of positive and negative correlation. For the cross-dataset alignment summaries, we also calculated the correlation separately within each dataset in the same draw and reported the fraction of draws in which all eight correlations shared a sign. The draws incorporated finite-cell feature uncertainty, spectral estimation uncertainty where applicable, and Bayesian-bootstrap uncertainty across patients.

The correlation figures show the posterior median for each feature pair. The clinical-field analysis used the arithmetic mean of the draw-specific correlation matrices.

### Clinical-field alignment across predictor regimes

#### Defining the clinical field and assigning regime weights

We constructed separate clinical association fields for TCAT GEP usages and atlas cell type proportions. For coordinate *j*, we defined

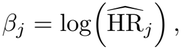

where HR_*j*_ was the population-level posterior median hazard ratio per global standard deviation increase in the CLR-transformed feature. Each clinical coefficient came from a separate continuous survival model adjusted for treatment and disease and accounting for dataset heterogeneity. We used the same coefficient vector in both regimes of each comparison.

We evaluated *O*_*λ*_ and the six cellular predictors whose adjusted smooth transition HRs had 95% credible intervals excluding one. Five were GEPs (CD8 naive, exhaustion, heat shock, activated Th17, and Tph), and the sixth was the antigen-specific activation composite score. We evaluated each context on both the GEP and atlas cell type geometries. Each context used its own fitted transition to define regime memberships. For each patient, *q*_*i*_ supplied the high-regime weight and 1− *q*_*i*_ the low-regime weight (Methods). The adverse regime was low for CD8 naive and high for the other six contexts.

When a context was itself a coordinate of the evaluated representation, we removed that coordinate and its coefficient. For ASA, we removed its four constituent GEPs: CTLA4/CD38, ICOS/CD38, OX40/EBI3, and TIMD4/TIM3. The ASA comparison therefore used 48 TCAT coordinates. Atlas composition thus remained unchanged for TCAT-defined contexts.

#### Estimating feature correlation matrices within each regime

For each of *B* = 500 realizations, we sampled one joint feature particle per patient and drew patient weights **v**^(*b*)^ ∼ Dirichlet(**1**). Joint sampling preserved dependence among features within each patient’s particle. For dataset *d* and feature *j*, we subtracted the weighted dataset mean

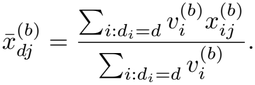

We then calculated weighted Pearson correlations of the dataset-centered profiles using weights proportional to 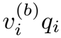 in the high regime and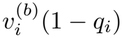 in the low regime. Dataset centering preceded regime weighting.

The Pearson calculation subsequently centered and standardized features under the weights of each regime. We defined the regime correlation matrix by the arithmetic mean

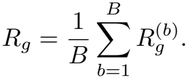

Arithmetic averaging preserves positive semidefiniteness. We first averaged the draw-specific matrices to obtain *R*_*g*_, then calculated each alignment statistic on *R*_*g*_. Positive-semidefiniteness diagnostics are reported in Supplementary analysis.

Dataset centering defined the adjustment for feature correlations. The clinical coefficients and regime memberships came from treatment- and disease-adjusted survival models and were held fixed across correlation draws.

#### Three measures of clinical-field alignment

For a nonzero clinical field ***β***, prognostic channeling was defined as

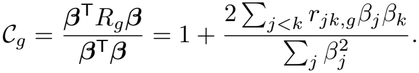

Channeling increases when positively correlated features have clinical associations of the same sign, or negatively correlated features have associations of opposing signs. Its reference value is one for an identity correlation matrix. This is an algebraic reference, rather than an independence null that must be attainable for a complete CLR composition. Because features are standardized separately within each regime, channeling measures alignment in correlation coordinates, not the variance of a common score on the original CLR scale.

For signed graph variation and signed-neighbor cosine, we used all off-diagonal pairs. Within each regime, let

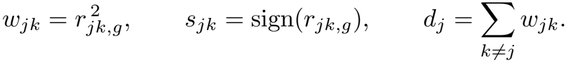

Signed graph variation was defined as

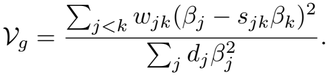

Lower values indicate closer agreement between the clinical associations of strongly correlated pairs, accounting for the sign of their correlation.

Finally, for *d*_*j*_ *>* 0, we defined the signed-neighbor field

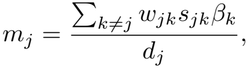

and defined signed-neighbor cosine as

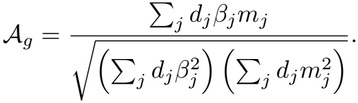

Higher values indicate closer agreement between the clinical field and the orientation implied by correlated neighbors. These feature graphs represent statistical relationships across patients, not physical interactions among programs or cell types.

All three statistics measure alignment with the clinical axis. Reversing the axis leaves their values unchanged, so they describe the pattern of variation within a regime independently of its average position along that axis. Contrasts were calculated as the adverse regime minus the favorable regime.

#### Within-dataset permutation tests of regime contrasts

For each context, we permuted soft membership weights within datasets 10,000 times. Each permutation retained the clinical field, feature particles, and patient-bootstrap draws, and recomputed both regime correlation matrices and all three contrasts. For metric *k*, we standardized the observed contrast against its permutation distribution,

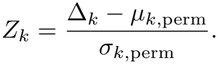

The joint statistic was then defined as

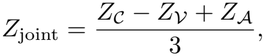

oriented so that greater channeling, lower variation, and stronger neighbor alignment in the adverse regime each increased the joint statistic. Applying the same transformation to each permutation retained dependence among the three metrics.

We calculated upper and lower-tail empirical probabilities with a plus-one correction:

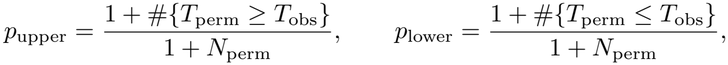

where *T* denotes an oriented individual statistic or the joint statistic. For the pooled analysis, we reported two-sided probabilities as *p* = min {1, 2 min(*p*_upper_, *p*_lower_)} . Within each representation, we applied the Benjamini–Hochberg procedure across the seven contexts, separately for the joint test and each individual metric. For comparison, the directional pooled joint probabilities for *O*_*λ*_ were 0.0004 (TCAT) and 0.0007 (atlas), corresponding to the two-sided probabilities reported in the Results.

This primary analysis tested whether alignment between cellular feature correlations and clinical risk differed across regimes. The held-out analysis tested whether this contrast generalized to new cohorts, and the surrogate-field tests assessed whether it was specific to the clinical direction estimated from patient outcomes.

### Pseudobulk gene associations and gene-set enrichment

We restricted expression analysis to the intersection of STRING-harmonized genes available across the included datasets. Let *y*_*ijg*_ denote the count for gene *g* in cell *j* from patient *i*, and let *n*_*i*_ be the number of retained infusion cells from that patient. For pseudobulk particle *b*, we drew cell weights

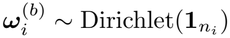

and calculated the weighted patient-level count

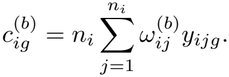

Within the shared gene universe *G*, each particle was transformed as

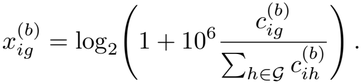

We generated 16 pseudobulk particles per patient, matching the particle representation used elsewhere in the uncertainty-propagation framework. Genes with constant or nonexpression across the analysis cohort were excluded. Particle construction was independent of clinical outcomes.

We estimated Bayesian-bootstrap partial Spearman correlations after adjustment for dataset. For each of 4,000 association draws, we selected one pseudobulk particle index, yielding one independently bootstrapped expression realization for every patient. We independently sampled one equal-probability *O*_*λ*_ particle per patient and reranked the sampled spectral estimates. We then generated patient weights

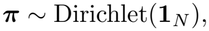

where *N* is the number of patients. Gene expression ranks and spectral ranks were separately regressed on an intercept and dataset indicators using these weights. The weighted Pearson correlation between the two residual vectors gave the draw-specific partial rank correlation *ρ*_*g*_. This procedure propagated finite-cell uncertainty in pseudobulk expression, uncertainty in the inferred spectral asymptote, and Bayesian-bootstrap uncertainty across patients.

For every gene, we summarized the resulting correlation distribution and ranked genes by

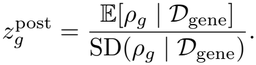

Here *D*_gene_ denotes the pseudobulk, spectral, and dataset information used in the gene-association analysis. The complete eligible ranking entered preranked enrichment; genes were not filtered by association magnitude or posterior directional probability. Gene symbols were used to order tied scores deterministically.

We used human Hallmark and Reactome gene sets from MSigDB version 2026.1.Hs [12], obtained through `msigdbr`. Gene sets containing 15–500 represented genes were tested using `fgseaMultilevel`, with `gseaParam=1`, `eps=0`, and `nPermSimple=1,000,000`. We applied Benjamini–Hochberg correction separately within each collection and used the resulting *q <* 0.05 criterion for the primary enrichment results. We also retained a correction across both collections. For every tested gene set, the output included its enrichment score, normalized enrichment score, leading-edge genes, nominal *p* value, within-collection FDR, and combined-collection FDR.

### Grouping Reactome pathways by shared genes

We constructed a pathway-overlap graph from all tested Reactome pathways with finite NES. For unique MSigDB gene sets *A* and *B*, similarity was computed as

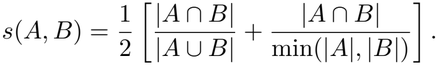

We retained edges with *s*(*A, B*) ≥ 0.45 and clustered the weighted graph by Markov clustering [13] with expansion order two and inflation 1.8. Similarity calculations used only pathway membership lists; NES, enrichment significance, and clinical outcomes entered neither similarity calculation nor clustering. The layout used positions derived from overlap topology and community membership, followed by similarity-weighted Fruchterman–Reingold refinement [14] with random seed zero. Community labels summarized official Reactome annotations and constituent pathway names. We fixed the pathway communities and layout using gene-set overlap before adding enrichment scores and significance to the network.

In the displayed network, color indicated NES, while node size and opacity emphasized pathway-level enrichment evidence only after the topology and community assignments had been fixed. Nonsignificant pathways were retained as context. The implementation displayed communities containing at least six pathways together with smaller communities connected to them through retained overlap edges. We labeled communities containing at least six pathways with Reactome *q <* 0.05. For each labeled community, we summarized the distribution of pathway NES values by its mean, full range, and 10th–90th percentile range.

## Code and Reproducibility

Analysis code is available at https://github.com/adamnik/CAR-T-spectral-organization. Dataset accessions are listed in Table 1.

## Supplementary analyses and information

### The complete-graph reference for diffusion lifetimes

For a connected, undirected, nonnegative weighted graph without self-loops, the symmetric normalized adjacency has zero trace:

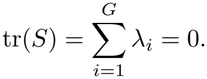

Removing the stationary eigenvalue *λ*_1_ = 1 gives the exact mean of the nonstationary eigenvalues,

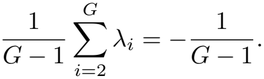

For an equal-weight complete graph on *G* vertices, every nonstationary eigenvalue equals this mean:

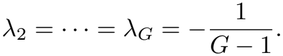

The corresponding reference lifetime is

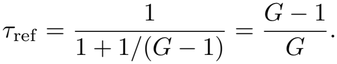

This quantity is the lifetime evaluated at the mean eigenvalue. Thus, dividing each modal lifetime by *τ*_ref_ ensures that every nonstationary mode of the equal-weight complete graph has relative lifetime one.

### Spectral organization approaches one at the equal-weight complete graph

Observe that *O*_*λ*_ → 1 as the graph approaches the equal-weight complete graph on *G* vertices. To see this, Let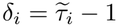. Then, whenever at least one *δ*_*i*_ is nonzero,

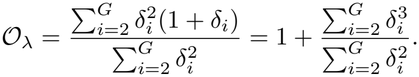

As the graph approaches complete uniformity, every nonstationary eigenvalue approaches 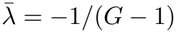, so 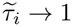 and *δ*_*i*_ → 0 for every *i* ≥ 2. Let 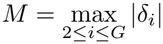. Then,

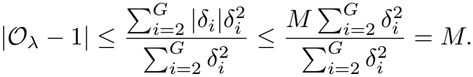

Since *M* →0, it follows that *O*_*λ*_ →1. At the equal-weight complete graph, both the numerator and denominator of the defining ratio vanish. The limit therefore gives a continuous extension of the statistic, and we define *O*_*λ*_ = 1 at this reference graph.

### Spectral organization is invariant to uniform edge-weight scaling

For a positive scalar *c*, let *W*^*′*^ = *cW* . Then *D*^*′*^ = *cD*, and

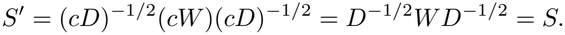

The spectrum, relative modal lifetimes, and *O*_*λ*_ are therefore unchanged. This identity concerns uniform scaling with the vertex set held fixed. It does not imply invariance to heterogeneous edge perturbations, gene selection, or finite-cell sampling.

### Empirical spectral domain and graph diagnostics

We audited 20,992 finite-cell trajectory graphs from the 163 patients retained in the primary analysis. The minimum was *O*_*λ*_ = 1.02653, obtained for Haradhvala patient 21 at 525 cells. All observations were finite and exceeded the log-excess fitting threshold of 1 + 10^*−*5^. These findings support the empirical domain used for asymptotic inference; they do not establish a universal lower bound of one for the statistic.

No audited graph required removal of isolated genes. Each graph had exactly one eigenvalue within 10^*−*5^ of unity, and the minimum nonstationary spectral gap, 1 − *λ*_2_, was 0.45099. The numerical safeguard bounding 1 − *λ*_*i*_ below by 10^*−*7^ was never activated. No graph triggered the near-uniform denominator rule assigning *O*_*λ*_ = 1. Thus, neither numerical safeguard affected the retained trajectory values.

### Validation of asymptote estimates from truncated trajectories

#### Derivation of the leading finite-depth correction

Fix a set of *G* genes, and let *W* denote their population mutual information matrix, with *W*_*gh*_ = *I*(*g, h*) for *g* ≠ *h* and *W*_*gg*_ = 0. Its estimate from *n* cells has off-diagonal entries 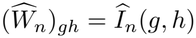. Assume that joint detection probabilities are interior, all weighted degrees are positive, and the population graph is connected. Write

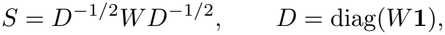

with eigenvalues 1 = *λ*_1_ *> λ*_2_ ≥ · · · ≥ *λ*_*G*_. Additionally, assume that 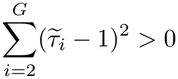.

For binary detection states, the plug-in mutual information estimator measured in nats satisfies

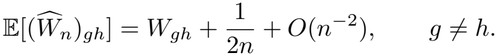

The estimation errors are correlated across edges because all pairs are computed from the same cells. Under the regular fixed-*G* expansion, write

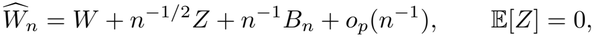

where *B*_*n*_ includes the second-order random terms.

Let *S*(*W* ) = *D*^*−*1*/*2^*WD*^*−*1*/*2^. Expanding the degree normalization gives

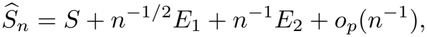

where

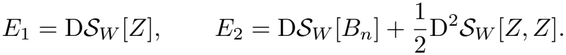

Thus, normalization contributes a second-order term even when the leading graph perturbation has mean zero.

For each simple nonstationary eigenvalue *λ*_*i*_, the expansion through order *n*^*−*1^ under a perturbation of *S* is

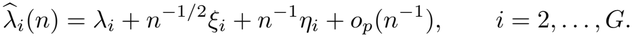

Here *ξ*_*i*_ and *η*_*i*_ are the first- and second-order perturbation coefficients for mode *i*, and ***ξ*** and ***η*** collect them across nonstationary modes. Let **g** and *H* denote the gradient and Hessian of *O*_*λ*_ with respect to the nonstationary eigenvalues, evaluated at the limiting spectrum. A second-order Taylor expansion then gives

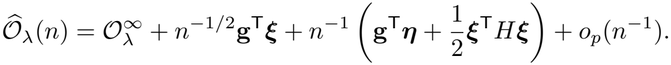

If E[***ξ***] = 0 and the remainder has expectation *o*(*n*^*−*1^), taking expectations gives the leading finite-depth bias,

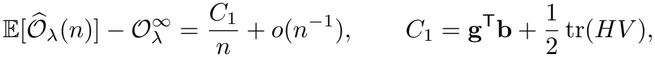

where **b** = E[***η***] and *V* = Cov(***ξ***). The sign of *C*_1_ depends on MI estimation, degree normalization, and spectral geometry.

When 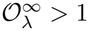, the same expansion on the log-excess scale again gives a leading bias term of order 1*/n*. To model this depth dependence, we use the bounded function

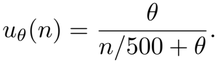

At large cell depths,

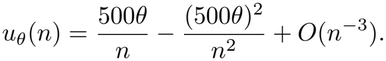

Thus, *b*_1_*u*_*θ*_(*n*) captures the leading 1*/n* correction, while *b*_2_*u*_*θ*_(*n*)^2^ allows the 1*/n*^2^ correction to vary independently. Both terms remain bounded at small depths and vanish as *n* → ∞.

#### Trajectory model and held-out patient validation

Finite-depth spectral organization was modeled on the log-excess scale,

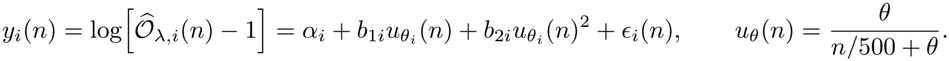

Because the finite-depth terms vanish as *n* → ∞,

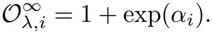

Unconstrained slope and curvature coefficients permitted both monotonic and nonmonotonic trajectories. A hierarchical population distribution informed patient-specific coefficients, and timescale uncertainty was integrated over nine equal-probability nodes under log *θ*_*i*_ = log 2 + 0.35*T*_*i*_, where *T*_*i*_ ∼ *t*_3_. Conditional slope deviations followed a bivariate Student-*t*_10_ prior, and patient-level updates used a Student-*t*_4_ likelihood with depth-dependent residual scale. Population parameters were estimated by maximum likelihood, followed by approximate empirical-Bayes patient updates and Laplace weighting across timescale components.

Because observations arose from nested cell subsets, the summed trajectory log likelihood was multiplied by min(1, 12*/m*_*i*_), where *m*_*i*_ was the number of fitting points. This capped the nominal information contribution at 12 effective observations without explicitly modeling covariance between depths. Within each timescale component, the inverse-Hessian scale of the fitted log-excess asymptote determined its Student-*t*_4_ uncertainty. We assessed the resulting intervals directly against full-trajectory estimates.

Validation included 16 patients with at least 7,500 available infusion product cells spanning the Li, Yu, Haradhvala, Cao, and Maurer datasets. For each patient, the population hierarchy was refitted after excluding all observations from that patient. The held-out trajectory was then fitted using only points through 1,000, 3,000, or 5,000 cells. A separate fit used the complete available trajectory with the same hierarchy learned without that patient. Model settings were held fixed throughout. This design prevented the held-out patient’s complete trajectory from informing its population prior.

#### Criteria for evaluating truncated-trajectory fits

We compared each patient’s central asymptote estimate from a truncated trajectory with the estimate from their full trajectory using mean absolute error (MAE), root mean squared error (RMSE), and Spearman rank correlation. Reference containment was reported as the number of finite-depth 95% intervals containing the complete-fit central estimate out of 16 patients, alongside median interval width. Withheld-trajectory MAE measured agreement between the prefix-fitted curve and observed points beyond the largest depth used in fitting. Errors were averaged within each patient and then across patients. The complete trajectory fit provided the strongest available reference estimate of the latent asymptote as the true infinite-depth value is unobserved.

#### Accuracy and uncertainty across sampling depths

At 1,000 cells, asymptote MAE relative to the complete fit was 0.1245, and patient ordering was only weakly preserved (Spearman *ρ* = 0.31; Supplementary Table S1). All 16 reference estimates were contained in the finite-depth intervals, and the median interval width was 0.793. These wide intervals reflect the limited information about the asymptote at 1,000 cells.

At 3,000 cells, rank correlation increased to *ρ* = 0.80, and median interval width decreased to 0.126. MAE decreased to 0.1019, although RMSE increased to 0.2105 because a small number of trajectories retained large errors. At 5,000 cells, rank correlation reached *ρ* = 0.94, MAE decreased to 0.0620, and median interval width was 0.061. Median absolute error was 0.0157, indicating substantially smaller error for the typical patient. Excluding the single most extreme trajectory reduced the 5,000-cell MAE to 0.0368.

The largest remaining errors occurred in high-asymptote trajectories that rose after the truncation depth. In these patients, the early trajectory contained little information about the later rise. Withheld-trajectory MAE decreased from 0.0292 at 1,000 cells to 0.0169 at 3,000 cells and 0.0101 at 5,000 cells, demonstrating improved prediction of subsequently observed finite-depth behavior.

At 3,000 cells, 13 of 16 intervals contained the complete-trajectory estimate (exact binomial 95% interval 0.544–0.960). At 5,000 cells, 12 of 16 did (0.476–0.927). These fractions measure agreement with an inferred reference in 16 high-depth patients. Supplementary Fig. S1A–C shows examples of how asymptote estimates and their uncertainty evolved as more cells were included.

### Survival validation in held-out cohorts

We performed two complementary analyses to assess the cross-cohort robustness of the spectral organization survival association; leave-one-dataset-out refitting tested whether the association depended on any single cohort, while cross-fitted evaluation tested whether survival scores learned from other cohorts remained associated with outcomes in the omitted patients. Both analyses used the full 163 patient population from the primary analysis.

#### Leave-one-dataset-out refitting

For each dataset *d*, we excluded all of its patients and separately refit the continuous and smooth transition survival models for *O*_*λ*_ using the remaining seven datasets. Each fit retained the primary model’s interval-censored likelihood, dataset-specific baseline hazards, hierarchically pooled predictor effects, treatment and disease adjustments, and integration over 16 *O*_*λ*_ particles per patient.

We used the same centering, scaling, and transition rules as in the primary analysis, recalculating all data-dependent quantities from the training patients alone. Similar to the primary survival analysis, for the smooth transition model, the crossover prior spanned the training 10th–90th percentiles, and the transition width was fixed at 0.10 training standard deviations. As so, the omitted dataset contributed neither to model fitting nor to these preprocessing and transition choices.

The continuous association remained adverse in every seven-cohort refit (median HR 1.39–1.67, *P* (*β >* 0) = 0.969–0.994). Six of eight 95% credible intervals excluded one, with wider intervals when Li or Yu was omitted.

The median transition HR ranged from 1.67 to 2.15, with *P* (*β >* 0) = 0.933–0.989. Five intervals excluded one. The remaining three fits omitted Bai2024, Haradhvala, or Li. The crossover medians occupied a narrow range (1.181–1.251; Supplementary Table S2).

#### Construction of cross-fitted survival scores

After each training fit, we constructed risk-score particles for patients in the omitted dataset. Let 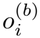 denote a particle representing patient *i*’s asymptotic *O*_*λ*_, and let *µ −*_*d*_ and *σ*_*−d*_ denote the mean and standard deviation of the central spectral estimates in the training population. For the continuous model, the score was

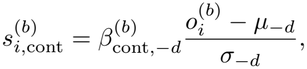

where 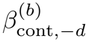 was drawn from the training posterior of the population-level coefficient.

For the smooth transition model, the score was

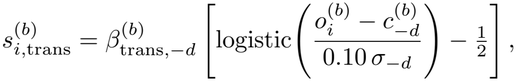

where 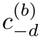 and 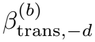 were paired draws from the training posterior. This preserved their posterior dependence. Patient spectral particles were randomly paired with the sampled training parameters to produce 16 equally weighted score particles per patient. Neither score used outcomes from the patient’s own dataset during construction. Furthermore, the clinical adjustment coefficients and baseline hazards from the training fits were not included in these scores; they represented the contribution of spectral organization alone.

#### Association of cross-fitted risk scores with treatment failure

Each patient contributed scores from the fold in which their complete dataset had been excluded. We then evaluated the continuous and transition scores separately using the hierarchical survival framework, fitting dataset-specific baseline hazards, partially pooled score coefficients, and treatment and disease adjustments against the combined held-out outcomes. Score particles were standardized using the global mean and standard deviation of their central patient estimates, and their uncertainty was marginalized in the likelihood.

**Supplementary Table S2:** Leave-one-dataset-out stability of the spectral organization survival association. Rows show posterior-median hazard ratios and 95% credible intervals from models fitted after excluding the indicated cohort and retaining the other seven datasets. Continuous HRs are reported per global standard deviation in the training population; transition HRs compare the high vs. low *O*_*λ*_ regimes. The final column gives the posterior-median crossover in native *O*_*λ*_ units. Results describe training-population stability. Held-out score evaluation is reported separately in Supplementary Table S3.

| Dataset omitted | Continuous HR [95% CrI] | Transition HR [95% CrI] | Crossover |
|---|---|---|---|
| Bai2022 | 1.43 [1.05, 2.13] | 2.01 [1.04, 3.44] | 1.206 |
| Bai2024 | 1.67 [1.07, 2.58] | 1.67 [0.82, 3.14] | 1.219 |
| Cao | 1.45 [1.10, 2.27] | 1.98 [1.10, 3.45] | 1.219 |
| Haradhvala | 1.41 [1.04, 2.12] | 1.82 [0.98, 3.27] | 1.224 |
| Li | 1.42 [0.96, 2.22] | 1.77 [0.90, 3.26] | 1.251 |
| Maurer | 1.39 [1.05, 2.07] | 2.08 [1.13, 3.60] | 1.198 |
| Wang | 1.42 [1.06, 2.19] | 1.94 [1.01, 3.55] | 1.207 |
| Yu | 1.46 [0.98, 2.39] | 2.15 [1.06, 3.98] | 1.181 |

**Supplementary Table S3:** Survival associations of dataset-cross-fitted spectral organization scores. Scores were constructed from survival models trained with the evaluated patient’s dataset excluded. HRs are reported per global standard deviation of each score. *P*_align_ gives the posterior probability that all dataset-specific validation effects share a direction. Evaluation re-estimated baseline hazards and clinical adjustments, testing score associations rather than frozen predictions of absolute risk.

| Cross-fitted score | HR [95% CrI] | *P*(*β* > 0) | *P*<sub>align</sub> |
|---|---|---|---|
| Continuous | 1.50 [1.04, 2.42] | 0.984 | 0.759 |
| Smooth transition | 1.84 [1.17, 2.93] | 0.996 | 0.705 |

Both cross-fitted scores were associated with increased treatment failure hazard. Per global standard deviation, the continuous score had an HR of 1.50 (95% CrI 1.04–2.42, *P* (*β >* 0) = 0.984) and the transition score an HR of 1.84 (1.17–2.93, *P* (*β >* 0) = 0.996). The corresponding probabilities of a common direction across all eight cohorts were 0.759 and 0.705 (Supplementary Table S3).

#### Sampling diagnostics and scope of survival validation

The seven-cohort training fits and the two cross-fitted score evaluations used two chains with 1,000 tuning iterations and 1,000 posterior draws per chain. The target acceptance probability was 0.99, and the random seed was zero. No divergent transitions were reported, and all reported maximum 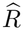 values were at most 1.01.

The validation assessed the survival association conditional on previously inferred spectral organization. Outcome-independent asymptote estimates and particles were carried into each fold without re-estimation. In addition, the combined held-out analysis estimated baseline hazards, clinical adjustments, and score coefficients anew. Therefore, these results regard the outcome association of cross-fitted scores, not the performance of a frozen model predicting absolute survival probabilities.

The adverse association persisted after each of the eight cohort omissions. Both continuous and transition scores remained prognostic in the pooled held-out patients when each patient’s score was learned from the other seven cohorts.

### Held-out validation and specificity of clinical-field alignment

#### Patient support for each spectral regime

Across the 163 patients, the low and high *O*_*λ*_ regimes had summed membership weights of 92.86 and 70.14, respectively. These totals describe how patient membership was divided between the two regimes. To assess how broadly each regime’s weight was distributed across patients, we also calculated its effective sample size,

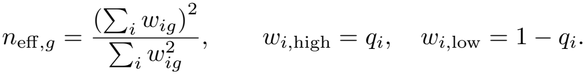

Using membership weights alone, the effective sample sizes were 118.22 for the low regime and 97.96 for the high regime, indicating that many patients contributed to each. Because patients could contribute to both regimes, these effective sample sizes can sum to more than 163.

#### Testing clinical-field alignment in held-out cohorts

For each held-out dataset, the other seven supplied both outcome-dependent quantities. We used their individual continuous survival coefficients for the GEP or atlas field and their midpoint-centered *O*_*λ*_ transition for the regime memberships. Feature scaling and transition width came from the same training cohorts. We averaged the training-derived activation over crossover draws and held-out patient spectral particles, then evaluated the training-derived field on the held-out feature correlations.

A cohort was evaluable when both summed membership weights, ∑_*i*_ *q*_*i*_ and ∑_*i*_(1− *q*_*i*_), were at least three. These quantities describe membership mass, not integer group sizes or effective sample sizes. Five cohorts met this criterion, comprising 144 patients (Supplementary Table S4). Cao, Maurer, and Wang did not meet the support criterion.

**Supplementary Table S4:** Support for held-out spectral regime comparisons. TCAT and atlas analyses used the same memberships, derived from transitions fitted with the evaluated dataset excluded. Comparisons required summed membership of at least three per regime; effective sample sizes were descriptive and were not used for eligibility.

|  |  | Membership sum |  | *n*<sub>eff</sub> |  |  |
|---|---|---|---|---|---|---|
| Dataset | *n* | Low | High | Low | High | Eligible |
| Bai2022 | 10 | 6.48 | 3.52 | 8.72 | 6.68 | Yes |
| Bai2024 | 42 | 22.18 | 19.82 | 27.39 | 25.18 | Yes |
| Cao | 9 | 6.63 | 2.37 | 8.25 | 5.27 | No |
| Haradhvala | 19 | 14.41 | 4.59 | 17.54 | 10.45 | Yes |
| Li | 35 | 23.97 | 11.03 | 28.79 | 17.34 | Yes |
| Maurer | 4 | 1.58 | 2.42 | 2.19 | 2.96 | No |
| Wang | 6 | 3.32 | 2.68 | 4.44 | 3.91 | No |
| Yu | 38 | 17.24 | 20.76 | 24.52 | 27.55 | Yes |

Within each eligible cohort, we generated 5,000 membership permutations and standardized each contrast against its cohort-specific null distribution. We combined oriented standardized contrasts with equal weight per cohort and evaluated their mean against the corresponding combined permutation distribution. Joint tests used the same orientation and three-metric average as the full cohort analysis.

Four of five cohorts had positive joint standardized scores in each representation. Haradhvala showed the opposite direction in both representations, and Bai2024 contributed the largest standardized atlas contrast. Each cohort received equal weight in the combined test.

For TCAT, the combined probabilities were 0.071 for channeling, 0.148 for signed graph variation, and 0.148 for signed-neighbor cosine, with joint *p* = 0.103. For atlas composition, the corresponding probabilities were 0.010, 0.021, and 0.013, with joint *p* = 0.011.

#### Correlation matrix diagnostics

We used arithmetic means of draw-specific correlation matrices throughout the pooled and held-out analyses. For every observed and membership-permuted mean matrix, we checked symmetry, the unit diagonal, and the minimum eigenvalue. We defined a positive-semidefiniteness failure as an eigenvalue below −10^*−*10^.

All 28 pooled and 20 held-out observed matrices passed these checks, as did the 280,000 pooled and 100,000 held-out permutation matrices. The smallest observed eigenvalues were 0.00323 in the pooled analysis and 0.00218 in the held-out analysis. Across permutations, the corresponding minima were 0.00274 and 0.00114. No positive-semidefiniteness correction was required.

#### Testing specificity to the clinical risk direction

While membership permutations ask whether spectral memberships distinguish alignment with a fixed clinical field, surrogate-field tests ask whether that contrast is unusually pronounced for the learned clinical direction, conditional on the observed regime geometries. We tested clinical-direction specificity within the compositional coordinates of each representation.

#### Compositional surrogate construction

TCAT usages and atlas proportions were analyzed as compositions. For a complete CLR representation with *p* coordinates, **1**^⊤^**x** = 0. Consequently, coefficient directions that differ only by a constant component define the same linear functional on CLR profiles. Feature standardization changes the direction of this constraint in correlation coordinates.

We therefore constructed surrogate fields in CLR coefficient coordinates and mapped both observed and surrogate fields through the same regime-specific transformations. Let *σ*_*j*_ be the global standard deviation used to scale feature *j* in its survival model. We first formed

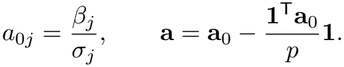

For held-out analyses, these scaling factors and clinical coefficients came from the training cohorts.

For draw *b* and regime *g*, let **s**_*gb*_ contain the feature standard deviations used in the weighted correlation calculation. The corresponding standardized CLR constraint is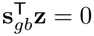. We defined

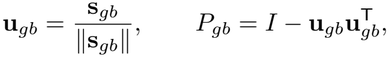

and averaged the draw-specific linear maps,

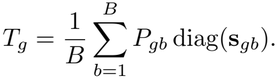

The sensitivity field in regime *g* was ***γ***_*g*_ = *T*_*g*_**a**. We evaluated the alignment statistics using the saved arithmetic-mean correlation matrix *R*_*g*_ and ***γ***_*g*_. Projection was performed within each draw before averaging the maps; the procedure did not assume that all draws shared one standardized constraint direction.

We generated 10,000 surrogate fields per family by permuting entries of **a**, independently randomizing signs of its absolute entries, or sampling Gaussian directions. For complete compositions, we removed each surrogate’s constant component and normalized it to ∥**a** ∥. Each surrogate was then mapped through the same pair of transformations, *T*_high_ and *T*_low_. Thus, observed and surrogate fields were compared under the same scaling and compositional constraints. For sign-randomized fields, feature-specific magnitudes were preserved before centering and normalization.

When one or more coordinates had been excluded, the retained original CLR coordinates no longer obeyed a zero-sum constraint among themselves. For these comparisons, we retained the global-to-regime standard deviation mapping but omitted centering and projection:

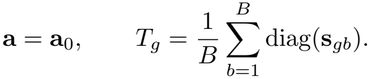

This rule applied to each defining GEP after its removal from the TCAT geometry (51 coordinates) and to ASA after removal of its four constituent GEPs (48 coordinates). We did not recompute CLR values for a new subcomposition.

Surrogate inference used the same directional standardization and joint statistic as the primary analysis. For pooled tests, Benjamini–Hochberg adjustment was performed across the seven contexts separately within each representation and surrogate family. Held-out probabilities were combined across the five eligible cohorts with equal weights and are reported as nominal probabilities. This sensitivity changes the scaling and projection of the observed field as well as the surrogate fields. It therefore provides a complementary test of clinical-direction specificity, rather than a repetition of the primary fixed-field membership test.

#### Surrogate-field results in pooled and held-out cohorts

We first compared high and low spectral organization regimes in the full cohort. For the TCAT analysis, we estimated the correlation geometry of 52 GEPs within each regime and defined the clinical field using the continuous-survival log hazard ratio for each GEP. The observed TCAT joint contrast exceeded all 10,000 coefficient-permuted and isotropic surrogates and all but one sign-randomized surrogate (*p* = 0.0001–0.0002, adjusted *p* = 0.0007–0.0014). The high spectral regime showed greater channeling (*p* = 0.0001–0.0002) and lower variation (*p* = 0.0001 for all three constructions). Signed-neighbor cosine provided weaker evidence of specificity (*p* = 0.072–0.094).

Using the same spectral regimes, we next estimated the correlation geometry of the 11 atlas phenotype proportions and defined the clinical field using the continuous-survival log hazard ratio for each proportion.

For atlas cell types, the pooled joint contrast had nominal support under all three constructions (*p* = 0.021– 0.034). After adjustment across contexts, the probabilities ranged from 0.148 to 0.236. Channeling provided the clearest individual-metric evidence (*p* = 0.0029–0.0148).

In a separate comparison, ASA defined the regimes. We excluded the four GEPs used to calculate ASA and evaluated the clinical field of the remaining 48 GEPs against their correlation geometry in the two ASA regimes. This comparison provided little evidence of joint specificity (*p* = 0.216–0.221).

For held-out validation, we returned to spectral organization regimes. For each evaluated cohort, the other cohorts supplied both the spectral transition used to assign regime memberships and the feature-specific survival coefficients defining the clinical field. The evaluated cohort supplied the GEP or atlas correlation geometry. We combined results across the five eligible cohorts with equal cohort weights (Supplementary Table S5).

For held-out GEP geometry, the joint probabilities ranged from 0.0276 to 0.0429. Channeling provided the clearest evidence across constructions (*p* = 0.0038–0.0062), with less consistent support for signed variation (*p* = 0.041–0.086) and weaker evidence for neighbor cosine (*p* = 0.229–0.246). For held-out atlas geometry, the joint probabilities ranged from 0.046 to 0.078. The corresponding ranges were 0.038–0.063 for channeling, 0.035–0.062 for variation, and 0.115–0.172 for neighbor cosine.

**Supplementary Table S5:** Joint compositional surrogate probabilities for spectral organization. Pooled and held-out analyses test clinical-direction specificity conditional on their respective regime geometries. Values are unadjusted.

| Representation | Surrogate family | Pooled | Held-out |
|---|---|---|---|
| TCAT | Coefficient permutation | 0.0001 | 0.0276 |
| TCAT | Sign randomization | 0.0002 | 0.0429 |
| TCAT | Isotropic directions | 0.0001 | 0.0297 |
| Atlas | Coefficient permutation | 0.0292 | 0.0463 |
| Atlas | Sign randomization | 0.0337 | 0.0784 |
| Atlas | Isotropic directions | 0.0212 | 0.0457 |

#### Interpretation

The alignment contrast is a property of the regime as a whole. An individual product can fall in the high *O*_*λ*_ regime with either a favorable or an adverse feature profile; the distinction lies in how features varied together across products in the high spectral regime. Taken together, these results suggest that the clinical relevance of gene-dependency architecture is also apparent in how cellular features collectively track risk across patients.

### Hallmark enrichment and Reactome pathway communities

Of the 50 tested Hallmark sets, 22 were positively enriched and nine were negatively enriched at within-collection *q <* 0.05 (Fig. 6). Of the 1,131 tested Reactome pathways, 184 were positively enriched and 202 were negatively enriched. These results were derived from the complete gene ranking; individual genes were retained regardless of their posterior directional probability.

Correction across Hallmark and Reactome together identified 419 gene sets at *q <* 0.05. All 417 sets meeting the within-collection criterion also met the combined threshold. Complete pathway statistics and leading edge gene lists are provided in the accompanying supplementary tables.

The Reactome overlap graph contained 1,131 nodes and 8,973 edges and yielded 179 communities. For visualization, the implementation retained communities containing at least six pathways and smaller communities reachable from them through retained overlap edges. This produced 1,043 displayed pathways. Sixteen communities met the additional labeling rule of at least six pathways with Reactome *q <* 0.05. Both rules were applied after clustering: the first controlled the displayed graph, and the second controlled community labels. All pathways assigned to each labeled community contributed to its NES summary.

The translation and ribosome-biogenesis community contained 16 pathways, all with negative NES, with mean NES = −2.86 and a range from −3.58 to −1.52. DNA replication and repair had negative NES for 40 of 43 pathways, with mean NES = −2.20. Nuclear transport and RNA processing and RNA polymerase II/TP53-associated transcription also had negative community means (−1.74 and −1.42, respectively), while containing individual pathways in both directions. Broader cell-cycle/APC-C and mitotic-checkpoint/Golgitrafficking communities had means near zero or weakly positive (0.12 and 0.26), despite containing strongly negative pathways.

Several communities associated with *O*_*λ*_ were uniformly positive. These included toll-like-receptor and innate-immune signaling (20 pathways, mean NES = 1.37), interleukin signaling (18 pathways, 1.41), complement, phagocytosis, and immune interactions (17 pathways, 1.57), and extracellular-matrix and collagen organization (19 pathways, 1.60). The glycosaminoglycan metabolism and biosynthesis community had a positive mean (1.27), with positive NES for 24 of 25 pathways. Receptor tyrosine kinase/PI3K– MAPK signaling and Rho-family GTPase communities also had positive means (1.06 and 1.05) but included pathways with negative NES. Uniformity of direction is distinct from the number of pathways meeting the FDR threshold.

### Cell resampling preserves the survival association, cellular features only partly reconstruct spectral organization

We evaluated whether spectral organization and its clinical association were sensitive to cell sampling, whether composition-matched cells from other patients reproduced the native spectral phenotype, and how much spectral organization could be reconstructed from established cellular-state representations. These analyses used patient-cell Bayesian bootstrap replicates, pooled-synthetic products matched by atlas-defined cell types, and patient-held-out reconstruction from TCAT GEPs and atlas cell type proportions.

#### Patient-cell resampling and graph construction

For each of 163 patients across eight datasets, we generated 25 posterior-predictive bootstraps containing 10,000 cells. Within each replicate, we drew cell weights

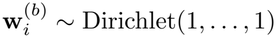

over the patient’s observed infusion cells and sampled with replacement according to these weights. The target resampling depth was therefore common across patients, although the number of distinct observed cells supporting each resample remained patient specific. Repeated draws of an observed cell contributed no new biological observations.

The eligible gene set was determined from each patient’s complete observed sample using the same two-state detection criterion as the trajectory analysis and held fixed across replicates. For every resample, we recomputed the binary mutual information graph and its spectral organization. We found that within-patient estimates were stable; the median replicate standard deviation was 0.0043, and the median width between the 2.5th and 97.5th replicate percentiles was 0.0146. Across patients, mean bootstrap estimates were concordant with the asymptotic estimates (Spearman *ρ* = 0.75).

#### Bootstrap spectral organization remains associated with treatment failure in the continuous model

We propagated all 25 replicate values per patient as equally weighted uncertainty particles through the continuous and smooth transition survival models, retaining the dataset hierarchy and clinical adjustments used in the primary analysis. The bootstrap analysis included the full 163 patients and 105 observed events.

The continuous model gave an HR of 1.51 (95% CrI 1.12–2.08, *P* (*β >* 0) = 0.994) for the patient-bootstrap estimates, compared with 1.45 (1.07–2.19, *P* (*β >* 0) = 0.989) for the asymptotic estimates. The posterior probability of a common direction across all eight datasets was 0.855 and 0.800, respectively. Despite the narrower between-patient spread of the finite-cell estimates (interquartile range 0.049 versus 0.186), the bootstrap transition model still yielded a high-versus-low HR of 1.80 (95% CrI 0.97–3.19, *P* (*β >* 0) = 0.969). The corresponding asymptotic HR was 2.03 (1.10–3.53). Across-dataset alignment was 0.741 and 0.800, respectively (Supplementary Table S6). Direct resampling preserved the continuous association and the adverse direction of the transition contrast. We retain 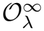 as the primary estimand because it targets a common infinite-depth limit, whereas resampling to a common depth still draws on different numbers of observed cells across patients. The narrow replicate intervals describe uncertainty within that observed support and do not replace uncertainty about the latent asymptote.

#### Synthetic products matched to each patient’s cell-type composition

For each target patient, we constructed synthetic products from the remaining patients in the same dataset, matching the target’s native atlas-defined cell type proportions through integer cell quotas. Within each cell type, quotas were divided approximately equally among eligible donors, and cells were sampled with replacement using independently drawn Dirichlet weights within donor–cell type blocks. Each synthetic product contained 10,000 cells. The target patient’s eligible gene set was retained, making this a compositional reference conditional on native gene eligibility.

**Supplementary Table S6:**
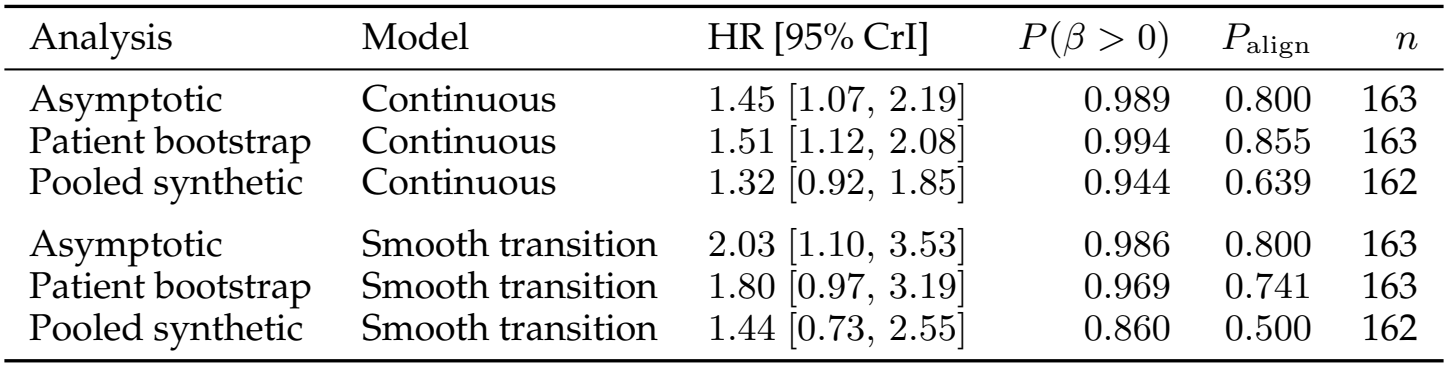
Survival associations under alternative estimates of spectral organization. Values are posterior medians and 95% credible intervals. Continuous HRs describe a global standard deviation increase, whereas smooth transition HRs compare the limiting high and low hazards in the midpoint-centered model. *P* (*β >* 0) denotes the posterior probability of a positive population-level coefficient; *P*_align_ denotes the probability that all dataset effects share a direction. The two resampling analyses used 25 uncertainty particles per patient.

Twenty-five replicates were available for each of 162 patients across all eight datasets. Composition matching was close: the largest summed absolute difference between target and realized cell type proportions was less than 0.00036. Synthetic estimates showed partial correspondence with both asymptotic spectral organization (*ρ* = 0.53) and patient-bootstrap estimates (*ρ* = 0.55).

For composition-matched synthetic products, the posterior-median HRs remained adverse but were smaller than for the patient-bootstrap estimates: 1.32 for the continuous model (95% CrI 0.92–1.85, *P* (*β >* 0) = 0.944) and 1.44 for the transition model (0.73–2.55, *P* (*β >* 0) = 0.860). All four sensitivity survival fits reported no divergent transitions. The synthetic analysis thus indicates that cohort context and coarse cellular composition reproduce part of the spectral signal, while substantial differences remain between native and substituted products.

#### Patient-held-out reconstruction from cellular states

We next evaluated reconstruction of asymptotic and mean bootstrap spectral organization from TCAT GEPs, atlas cell type proportions, and both representations together. We compared ridge regression, elastic net, partial least squares (PLS), and shallow gradient boosting using the same three repeats of four dataset-stratified patient folds for both targets. Each dataset contributed patients to training and testing. Within training data, three-fold validation selected hyperparameters by mean absolute error on the original lifetime scale. Dataset-specific feature and target means and feature scaling were learned anew in each training split. We restored training-derived dataset target means before averaging the three held-out predictions for each patient.

We calculated pooled *R*^2^ on the original scale. For dataset-adjusted Spearman *ρ*, we ranked predictions and observations across patients, subtracted the corresponding dataset means from both rank vectors, and correlated the residual ranks. Intervals used 1,000 within-dataset patient resamples of the saved predictions. The targets were the central asymptotic estimate and the mean bootstrap estimate per patient. A dataset-mean predictor was evaluated separately as a baseline.

Cellular states recovered a meaningful but incomplete component of spectral organization (Supplementary Table S7). For 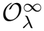, ridge regression on the TCAT GEPs achieved the highest observed reconstruction (*R*^2^ = 0.313, 95% bootstrap interval 0.199–0.390, dataset-adjusted *ρ* = 0.42). Atlas proportions alone achieved *R*^2^ = 0.216 with gradient boosting and the combined representation achieved *R*^2^ = 0.294 with PLS. GEP-only *R*^2^ ranged from 0.179 to 0.313 across the four model classes. Adding atlas proportions did not improve the best observed reconstruction.

The best reconstruction of the patient-bootstrap target also used GEPs (PLS *R*^2^ = 0.380, 95% bootstrap interval 0.154–0.515, dataset-adjusted *ρ* = 0.47). Atlas-only and combined models achieved *R*^2^ = 0.147 and 0.343, respectively. Dataset means alone yielded *R*^2^ = 0.018 for the asymptotic target and − 0.072 for the bootstrap target. In summary, even the highest-performing models left substantial variation unreconstructed, particularly for the asymptotic estimates.

#### Interpretation

Together, these analyses support the robustness of spectral organization while defining its relationship to known cellular states. Bootstrap estimation preserved the continuous association with treatment failure and yielded a transition contrast in the same adverse direction. Furthermore, composition-matched synthetic products and multivariate cellular-state models reproduced part of the signal, but neither convincing recovered the complete patient-level spectral phenotype.

**Supplementary Table S7:**
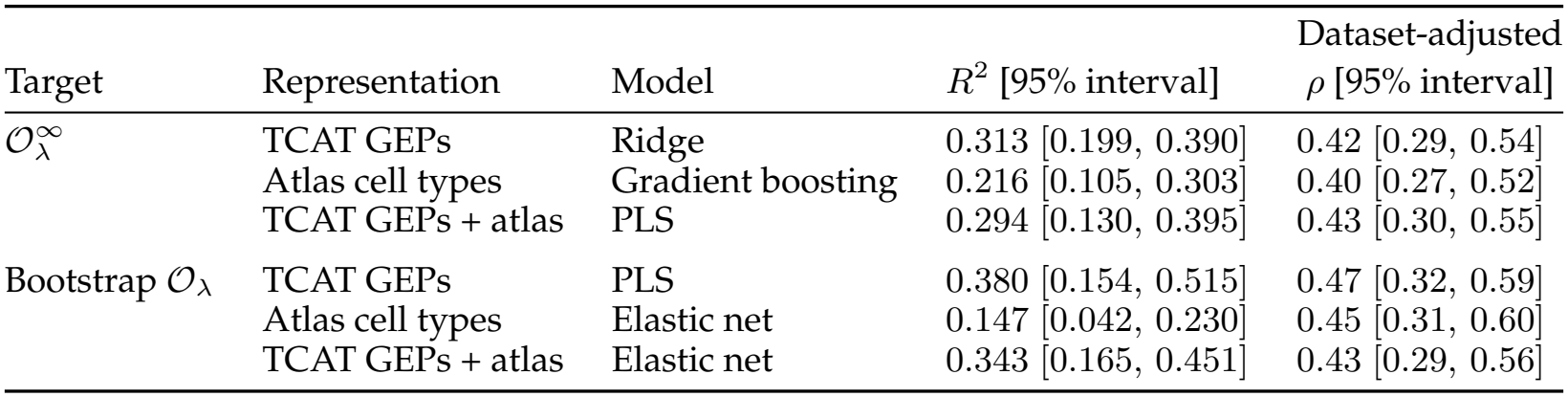
Partial reconstruction of spectral organization from cellular-state representations. Results are shown for the model class with the highest observed cross-validated *R*^2^ within each representation and target among ridge regression, elastic net, PLS, and shallow gradient boosting. Intervals are 95% dataset-stratified patient-bootstrap intervals conditional on the saved predictions; they do not incorporate full model-refitting or model-class selection uncertainty. The pooled *R*^2^ uses predictions on the original lifetime scale after restoring training-derived dataset offsets and averaging held-out predictions across repeats. Spearman *ρ* is adjusted for dataset by centering both rank vectors within datasets.

|  |  |  |  | Dataset-adjusted |
|---|---|---|---|---|
| Target | Representation | Model | *R*<sup>2</sup> [95% interval] | ρ [95% interval] |
| 𝒪<sub>λ</sub><sup>∞</sup> | TCAT GEPs | Ridge | 0.313 [0.199, 0.390] | 0.42 [0.29, 0.54] |
|  | Atlas cell types | Gradient boosting | 0.216 [0.105, 0.303] | 0.40 [0.27, 0.52] |
|  | TCAT GEPs + atlas | PLS | 0.294 [0.130, 0.395] | 0.43 [0.30, 0.55] |
| Bootstrap 𝒪<sub>λ</sub> | TCAT GEPs | PLS | 0.380 [0.154, 0.515] | 0.47 [0.32, 0.59] |
|  | Atlas cell types | Elastic net | 0.147 [0.042, 0.230] | 0.45 [0.31, 0.60] |
|  | TCAT GEPs + atlas | Elastic net | 0.343 [0.165, 0.451] | 0.43 [0.29, 0.56] |

### Technical and cellular covariates explain limited variation in spectral organization

We asked whether spectral organization could be reconstructed from sampling depth, sequencing quality, basic graph properties, or simple summaries of cellular heterogeneity. Across the full 163 patients, we evaluated ten covariates using both the asymptotic estimate 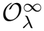 and bootstrap estimates of *O*_*λ*_. These included retained cell number, counts and detected genes per cell, mitochondrial fraction, graph node count, graph density, mean positive mutual-information weight, cell type composition entropy, effective rank of the cell-state covariance, and mean pairwise cell-state distance. Bayesian regressions incorporated dataset heterogeneity and patient-specific uncertainty in the spectral estimates and cell-derived covariates.

#### Variance explained by the joint covariate model

When all ten covariates were included together, the posterior median explained variance was *R*^2^ = 0.096 for 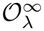 (95% CrI 0.015–0.222) and *R*^2^ = 0.167 for bootstrap *O*_*λ*_ (95% CrI 0.067–0.289; Supplementary Table S8). In other words, the combined measurements captured only 9.6% and 16.7% of modeled variation, respectively. Most spectral variation remained unexplained even after accounting jointly for conventional technical measurements, coarse properties of the inferred graph, and summaries of product heterogeneity. These estimates describe reconstruction under the fitted joint regressions.

#### Sequencing quality and cell count show weak associations with spectral organization

Retained cell number, counts per cell, mitochondrial fraction, and detected genes per cell all had global-effect intervals spanning zero for both 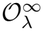 and bootstrap *O*_*λ*_. For the asymptotic estimate, cell number and mitochondrial fraction each had median individual *R*^2^ *<* 0.02, while counts and detected genes per cell had median *R*^2^ of approximately 0.06. Their contributions to bootstrap *O*_*λ*_ were similarly small, with median *R*^2^ values ranging from 0.007 to 0.036. Thus, given the agreement between the two spectral estimates, neither sampling depth nor standard sequencing quality metrics readily explain the spectral differences among infusion products.

Basic edge summaries also captured little spectral variation. Graph density and mean positive mutual information weight had median individual *R*^2^ values of 0.029 and 0.010, respectively, for 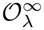, and 0.008 and 0.035 for bootstrap *O*_*λ*_. Their global effect intervals included zero in both analyses. Hence, products with similar overall graph density or average dependency strength could therefore differ substantially in spectral organization, consistent with the statistic describing how dependencies are distributed across graph modes.

#### The association with graph size varies across datasets

Graph node count showed the largest individual contribution to the asymptotic estimate, with median *R*^2^ = 0.228 (95% CrI 0.106–0.356). Its global standardized effect remained uncertain (0.251, 95% CrI − 0.098–0.587), with substantial variation in dataset-specific slopes (posterior-median dataset-effect SD = 0.385). The posterior probability that all datasets shared a direction was 0.188. For bootstrap *O*_*λ*_, graph node count captured considerably less variation, with median *R*^2^ = 0.022.

The individual graph-size contribution exceeds the joint-model estimate because the models represent dataset heterogeneity differently. Individual regressions allow dataset-specific slopes, whereas the joint model estimates shared coefficients with shrinkage. In the joint model, graph size retained a positive conditional association with asymptotic spectral organization (standardized coefficient 0.246, 95% CrI 0.010– 0.769). Even with graph size included, the shared covariates explained only a modest fraction of modeled variation.

**Supplementary Table S8:**
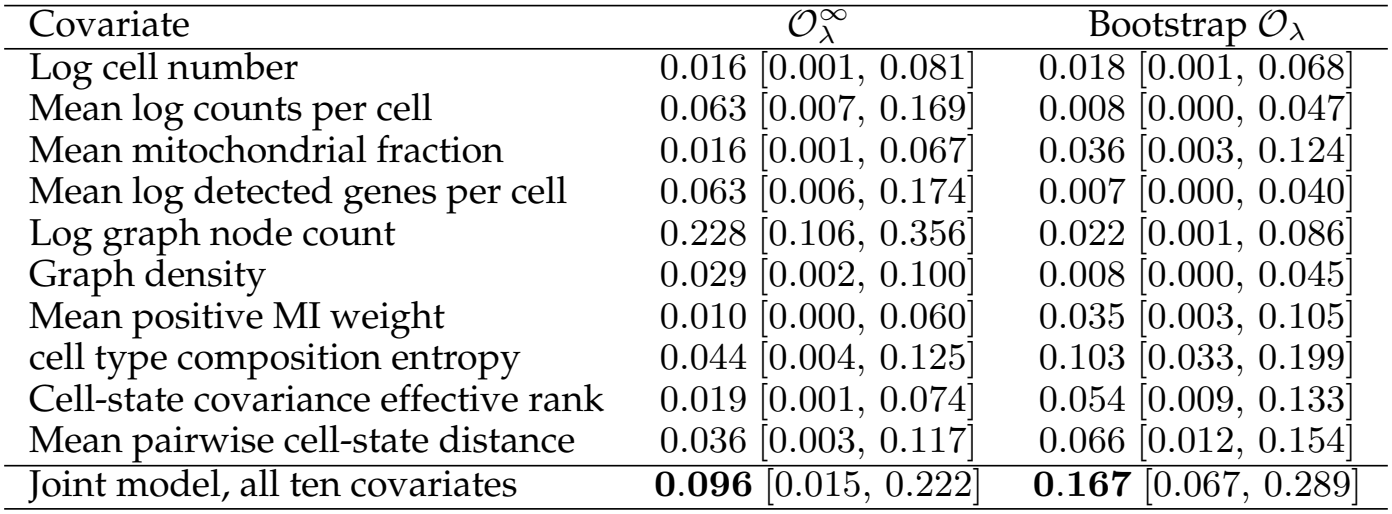
Limited reconstruction of spectral organization from technical and cellular-heterogeneity covariates. Values are posterior-median model-based *R*^2^ with 95% credible intervals. Individual regressions included hierarchical dataset intercepts and slopes. The joint model included all ten covariates with shared, regularized coefficients and hierarchical dataset intercepts. Explained variance was calculated from technical predictions using covariates centered within datasets, excluding variation attributable solely to dataset intercepts.

| Covariate | 𝒪<sub>λ</sub><sup>∞</sup> | Bootstrap 𝒪<sub>λ</sub> |
|---|---|---|
| Log cell number | 0.016 [0.001, 0.081] | 0.018 [0.001, 0.068] |
| Mean log counts per cell | 0.063 [0.007, 0.169] | 0.008 [0.000, 0.047] |
| Mean mitochondrial fraction | 0.016 [0.001, 0.067] | 0.036 [0.003, 0.124] |
| Mean log detected genes per cell | 0.063 [0.006, 0.174] | 0.007 [0.000, 0.040] |
| Log graph node count | 0.228 [0.106, 0.356] | 0.022 [0.001, 0.086] |
| Graph density | 0.029 [0.002, 0.100] | 0.008 [0.000, 0.045] |
| Mean positive MI weight | 0.010 [0.000, 0.060] | 0.035 [0.003, 0.105] |
| cell type composition entropy | 0.044 [0.004, 0.125] | 0.103 [0.033, 0.199] |
| Cell-state covariance effective rank | 0.019 [0.001, 0.074] | 0.054 [0.009, 0.133] |
| Mean pairwise cell-state distance | 0.036 [0.003, 0.117] | 0.066 [0.012, 0.154] |
| Joint model, all ten covariates | **0.096** [0.015, 0.222] | **0.167** [0.067, 0.289] |

#### Associations with cell type composition and cell state heterogeneity

For bootstrap *O*_*λ*_, cell type composition entropy had a positive standardized association (0.292, 95% CrI 0.028–0.556), while effective rank of the cell-state covariance had a negative association (−0.244, 95% CrI −0.463 to −0.008). Their median individual *R*^2^ values were 0.103 and 0.054, respectively. These measures describe different aspects of product heterogeneity, reflecting the evenness of cell type composition and the distribution of variation across embedding dimensions. Their associations with 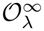 were weaker, with global-effect intervals spanning zero. Mean pairwise cell-state distance also had an interval spanning zero for both spectral estimates.

Collectively, the results show that spectral organization is poorly captured by the measured technical and cellular heterogeneity covariates. Conventional quality measurements showed consistently weak associations with spectral organization. Even with graph size included, the combined covariates left most modeled spectral variation unexplained. These findings argue against the measured technical factors as the primary explanation for differences among products.

#### Sampling diagnostics for the covariate models

All individual regressions and the bootstrap joint model had zero divergent transitions. The asymptotic joint model had one divergent transition. The largest reported 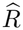 was 1.01, and the smallest bulk effective sample size was 1,115.

### Survival associations of spectral organization, spectral gap, and normalized Kemeny’s constant

We compared spectral organization with two simpler summaries of the same normalized adjacency spectrum: the spectral gap,

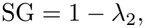

and the complete-graph-normalized Kemeny constant,

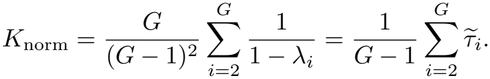

The spectral gap characterizes the slowest nonstationary mode, whereas the normalized Kemeny’s constant averages the relative lifetimes of all nonstationary modes equally. Spectral organization instead weights these lifetimes by their squared departure from the complete-graph reference.

For each of the 163 patients, we generated 25 graph replicates by drawing 10,000 cells with replacement using Bayesian-bootstrap cell weights, while retaining the patient’s native full-depth gene universe. The resulting metric values supplied equal-weight particles, with their arithmetic mean serving as the central estimate. These benchmarks therefore used finite-cell resampling rather than asymptotic trajectory inference.

Each metric was evaluated separately using the continuous and smooth transition survival models, with treatment and disease adjustment, dataset-specific baseline hazards, partially pooled associations, and likelihood integration over the 25 particles. Continuous hazard ratios were expressed per global standard deviation increase. Smooth transition models used crossover priors spanning the 10th–90th percentiles and midpoint-centered activations with a width of 0.10 global standard deviations (Methods).

In the continuous models, the spectral gap hazard ratio was 0.83 (95% CrI 0.61–1.13), and the normalized Kemeny hazard ratio was 0.84 (0.57–1.17). The corresponding high-versus-low transition HRs were 0.76 (0.40–1.40) for spectral gap and 0.69 (0.34–1.43) for normalized Kemeny. All four benchmark intervals included one. Among the three bootstrap statistics, only *O*_*λ*_ had a continuous interval excluding one (HR 1.51, 1.12–2.08). Its transition HR was 1.80 (0.97–3.19) (Supplementary Fig. S2). No divergent transitions were recorded, and reported 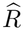 values were 1.00.

Thus, neither the slowest mode alone nor the equally weighted mean lifetime showed a clearly resolved outcome association in these analyses. At the same bootstrap depth, the continuous association with treatment failure was clearest when modes were weighted by their departure from the uniform reference. This identifies the distribution of persistence across those modes as a clinically relevant feature of the infusion-product graph.

**Supplementary Fig. S2:**
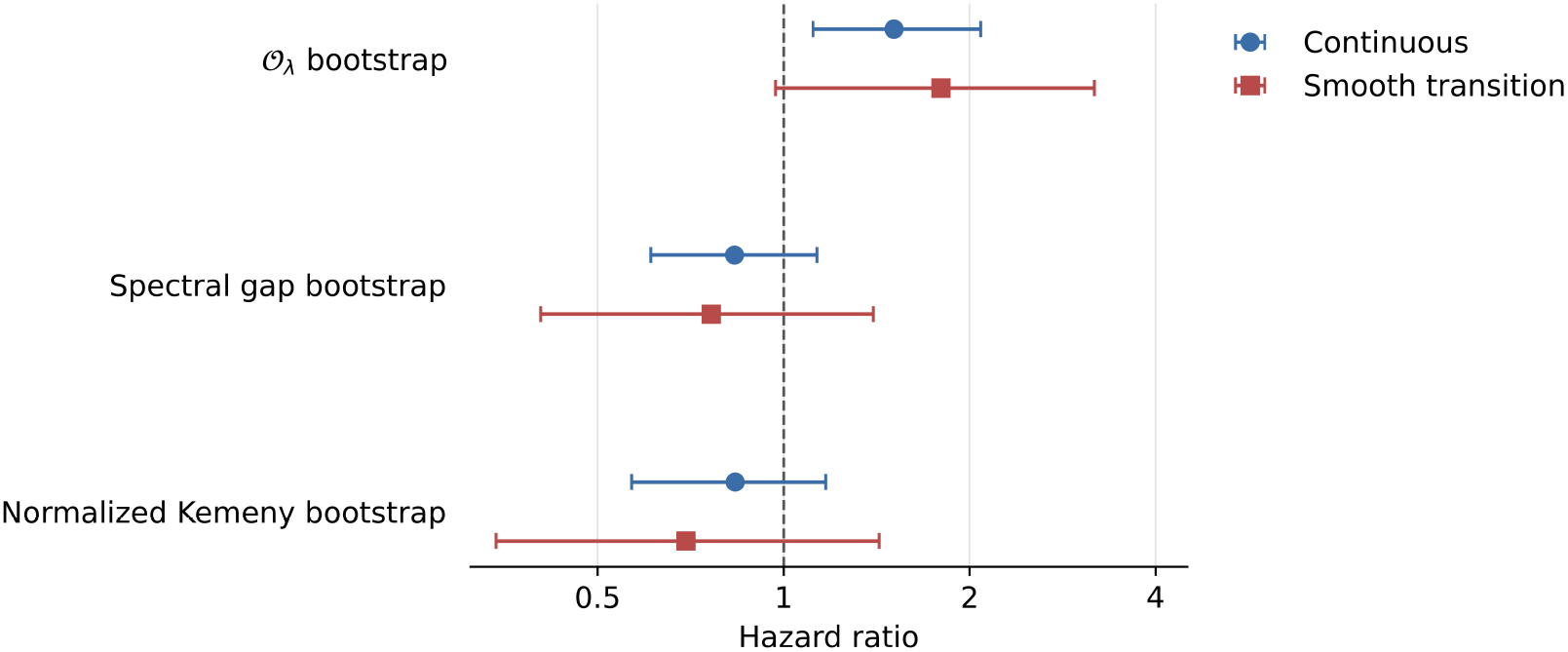
Survival associations of bootstrap spectral statistics. Posterior-median hazard ratios and 95% credible intervals are shown for patient-bootstrap estimates of spectral organization (*O*_*λ*_), spectral gap, and normalized Kemeny. Continuous effects are reported per global standard deviation, whereas smooth transition effects compare the limiting high and low hazards with the baseline at the transition midpoint. All models propagated uncertainty across patient-bootstrap graph realizations and included dataset-specific baseline hazards, hierarchically pooled dataset effects, and treatment and disease adjustments. Hazard ratios above one indicate greater treatment failure hazard.

## Footnotes

1 This model restricts the inferred asymptote to 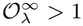. This modeling choice was motivated by the observation that all audited finite-depth values exceeded one (Supplementary diagnostics), rather than a universal bound on the statistic.

